# Propagated circulating tumor cells uncovers the role of NFκB and COP1 in metastasis

**DOI:** 10.1101/2022.12.09.519805

**Authors:** Jerry Xiao, Utsav Sharma, Abolfazl Arab, Sohit Miglani, Sonakshi Bhalla, Shravanthy Suguru, Robert Suter, Reetu Mukherji, Marc E. Lippman, Paula R. Pohlmann, Jay Zeck, John L. Marshall, Benjamin A. Weinberg, Aiwu Ruth He, Marcus S. Noel, Richard Schlegel, Hani Goodarzi, Seema Agarwal

## Abstract

Circulating tumor cells (CTCs), a population of cancer cells that represents the seeds of metastatic nodules, are a promising model system for studying metastasis. However, expansion of patient-derived CTCs *ex vivo* is challenging and dependent on the collection of high numbers of CTCs, which are ultra-rare. Here, we report the development of a combined CTC and CTC-derived xenograft (CDX) platform for expanding and studying patient-derived CTCs from metastatic colon, lung, and pancreatic cancers. Propagated CTCs yielded a highly aggressive population of cells that could be used to routinely and robustly establish primary tumors and metastatic lesions in CDXs. Differential gene analysis of the resultant CTC models emphasized a role for NF-kB signaling as a pan-cancer signaling pathway involved in metastasis. Furthermore, metastatic CTCs were identified through a prospective 5-gene signature (*BCAR1, COL1A1, IGSF3, RRAD, and TFPI2*). Whole-exome sequencing of CDX models and metastases further identified mutations in constitutive photomorphogenesis protein 1 (*COP1*) as a potential driver of metastasis. These findings illustrate the utility of the combined patient-derived CTC model and provide a glimpse of the promise of CTCs in identifying drivers of cancer metastasis.

## INTRODUCTION

As high as 90% of all cancer-associated deaths are directly attributable to the metastatic spread of cancer cells from the primary site to a secondary organ^1, 2^. Emphasizing its important role, metastasis was listed as one of the six hallmarks of cancer, which describe the biological capabilities acquired during the multistep development of tumors^3^. However, despite its role in cancer, metastasis remains a mysterious phenomenon largely due to a lack of clinically robust and accessible individualized patient-derived models^4, 5^.

Previous patient-derived models for identifying drivers of metastasis have depended on solid tissue biopsies of matched primary and metastatic lesions^6, 7^. While numerous prospective therapeutic compounds have successfully inhibited metastases in preclinical models, most have been unsuccessful in clinical studies^8^, suggesting that these models do not faithfully replicate clinical metastasis. For instance, early studies evaluating SRC inhibitors dasatinib and saracatinib in various cancer models (pancreatic, thyroid, prostate, ovarian, and melanoma) showed promise preclinically, but failed to demonstrate efficacy in human trials^8–10^. Thus, the frequent failure of these and other promising candidates in preventing metastasis in clinical trials suggests that these preclinical models—which often rely on either cell line-based or tumor explants—do not faithfully recapitulate clinical metastasis^8^.

An evolving approach to understanding metastases is to leverage circulating tumor cells (CTCs)^10^ which have detached from the primary tumor and/or from a metastatic site and entered the circulatory system, thereby undergoing the early stages of metastasis^11^. CTCs can be obtained from routine blood draws, subjecting patients to minimal discomfort in addition to providing easy access for serial monitoring of disease^11–15^.

Importantly, recent advances have demonstrated that CTCs are a highly heterogeneous, information-rich and a distinct metastatic driven rare population of cancer cells that may provide in-depth insights into metastasis^12, 13^. Simultaneously, it is not possible currently to fully capitalize on the considerable promise of CTCs due to challenges associated with identifying and obtaining CTCs in sufficient numbers to carry out mechanistic and functional studies^11^. The primary difficulty is that CTCs are ultra-rare, typically numbering < 10 cells per mL of blood^16^. As a result, molecular and mechanistic studies that exploit CTCs require first expanding CTCs *ex vivo*^11^. Such methods fall into two broad categories, namely, *in vitro* cell culture and *in vivo* CTC-derived xenografts (CDXs)^17, 18^. Unfortunately, these methods are also dependent on the ability to isolate relatively large quantities of CTCs to initiate cultures. On average, only 10-25% of all initiated CTC cultures result in successful cultures^11, 17, 19–23^. Similarly, the formation of patient-derived CDXs has been difficult to achieve when extracted CTC counts are lower than 400 CTCs/7.5 mL blood^18, 24–26^. Thus, CTC cultures and CDX models have been difficult to establish on most patients.

Once successfully propagated, mechanistic studies of patient-derived CTC cultures and CDXs have already contributed to key efforts in understanding metastatic phenomenon^11^. For instance, transcriptomic profiling of breast-cancer derived CTCs reaffirmed the importance of mTORC1, TGF-β, and p53 signaling in metastasis^27^. Another report demonstrated that CTCs from a patient diagnosed with estrogen receptor (ER) positive bilateral breast carcinoma and bone metastases could be cultured without estrogen supplementation, emphasizing a clinically relevant phenomenon of the decoupling of estrogen signaling and cell survival^28^. Treatment of CDX models has also been shown to accurately mirror the response of the patient to therapies^29^. For instance, CDX models derived from patients diagnosed with small cell lung cancer showed reduction in tumors following cisplatin and etoposide treatment, consistent with the response of the patients from whom the tumors were derived^30^.

By contrast, we recently published a novel method using a density-dependent, non-biased methodology (i.e., without the use of antibodies) for selecting and expanding CTCs and associated CD45^+^ cells in culture from patients with metastatic breast cancer^27^. Following our report, others have also confirmed the presence of co-inhabiting CD45^+^ cells in successful short-term pancreatic CTC cultures^31^. Here, we report the creation of patient-derived CTC cultures and CDX models from patients diagnosed with colon, lung, and pancreatic cancer. Notably, these CTC-derived CDX models were characterized by highly aggressive, de-differentiated primary tumors with an ability to form macro-metastatic lesions similar to their corresponding patient tumors. Pathway and differential gene expression analyses comparing patient-matched whole blood, cultured CTCs, and corresponding CDX models also highlighted a potential cancer-agnostic role for NF-kB signaling crucial to metastasis. Furthermore, differential gene analysis of metastatic CTCs pared down alterations in RNA expression amongst CTCs to reveal 5 candidate genes whose expression correlated with poorer prognosis in the TCGA pan-cancer dataset. Finally, whole-exome sequencing comparing mutations between formalin-fixed paraffin-embedded (FFPE) tissue biopsies, cultured CTCs, and CDX primary tumors identified both mutational concordance/discordance as well as a prospective driver of cancer metastasis in COP1. Taken together, the novel *in vitro* and *in vivo* models established in this study have contributed to the identification of key molecular signatures associated with cancer metastasis.

## RESULTS

### CTCs can be established from multiple cancer types

Previously, we reported the successful initiation of short-term cultures of breast-cancer derived CTCs using a novel methodology^27^. Here, we demonstrate that this platform can be applied to additional cancer types, namely, lung, colon, and pancreatic cancers. These cancers were selected as they are among the most common cancers diagnosed in the United States, and together accounted for over 270,000 deaths in 2021^32^. To begin, we obtained 25 liquid biopsies from 21 consented individual patients and processed these samples for expansion of CTCs as described previously (see reference 27 and **Fig. 1**)^27^. Short-term cultures with viable cells were established from 5/7 colon, 5/5 lung, and 13/13 pancreatic cancers, resulting in an overall success-rate of 92.0% for establishing CTC cultures. Thus far CTC cultures derived from lung cancers were observed to survive the longest in culture (59 ± 27 days), followed by colon (56 ± 25 days) and pancreatic (39 ± 20 days) cancers. qRT-PCR amplification confirmed expression of CTC markers Cytokeratin 18 (CK18) and Vimentin (VIM) in all cultures (**Fig. S1**).

**Figure 1:**
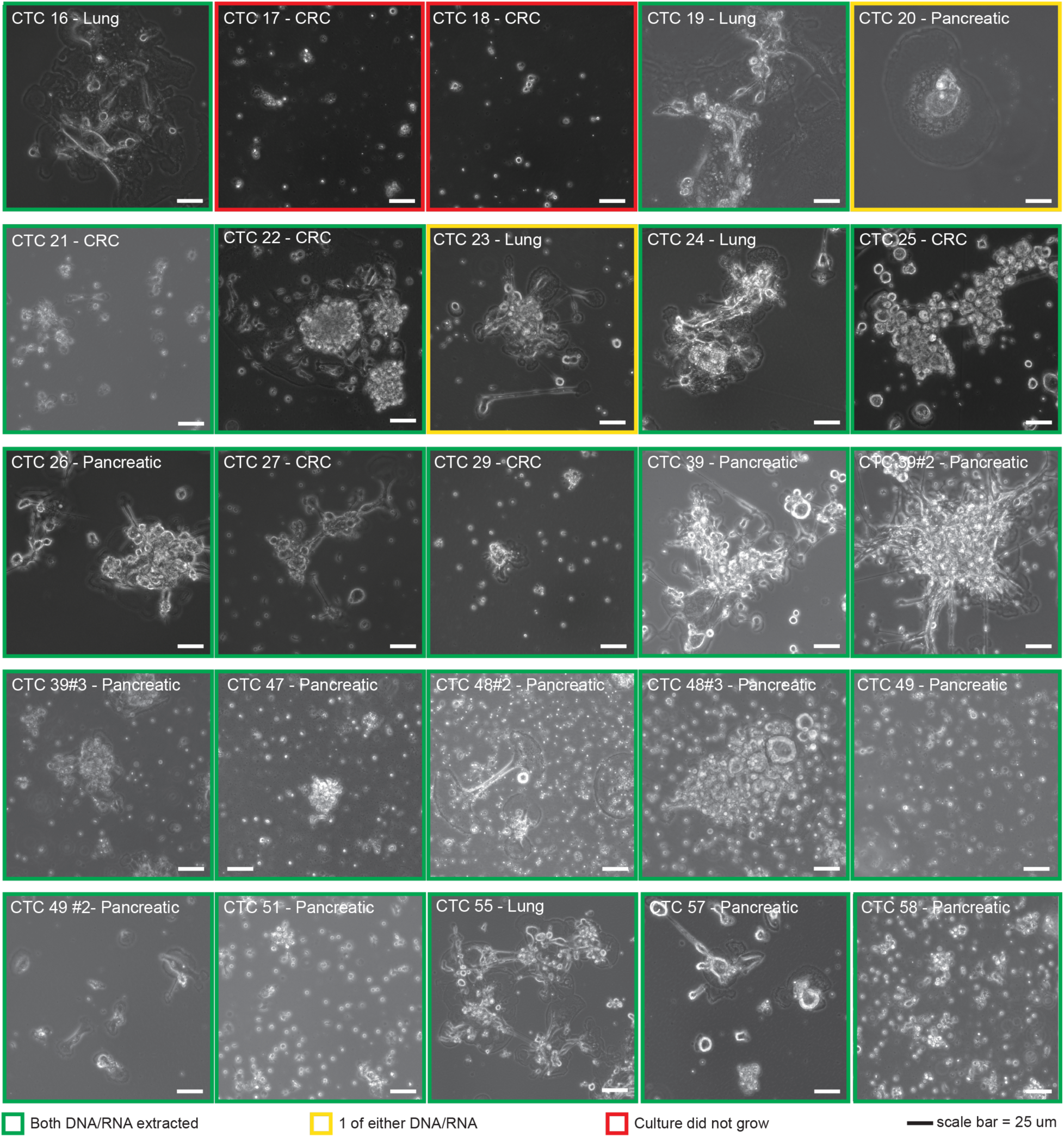
CTCs derived from multiple cancer types can be cultured. (a) Representative phase images of CTC cultures derived from lung, colon, and pancreatic primary tumors are shown. Images are outlined based on the ability to gather high-quality RNA or genomic DNA. Scale bar = 25 um.

### CDXs can be established from CTCs expanded in vitro

Previously, the single most important criterion for successfully establishing CDX models from liquid biopsy has been a high initial extracted CTC count (≥400 CTCs)^11, 18, 33–35^. Unfortunately, patients with a high CTC count in peripheral blood are extremely rare as usually <10 cells/ml are present in the patient’s blood. We therefore hypothesized that patient-derived CDX models could be more routinely established following an initial expansion phase in *in vitro* CTC cultures. Four patients with high-growth CTC cultures derived from lung (Patient #19), colon (Patients #21, & 22) and pancreatic (Patient #26) adenocarcinomas were selected as candidates for establishing CDX models (**Fig. 2a**).

**Figure 2:**
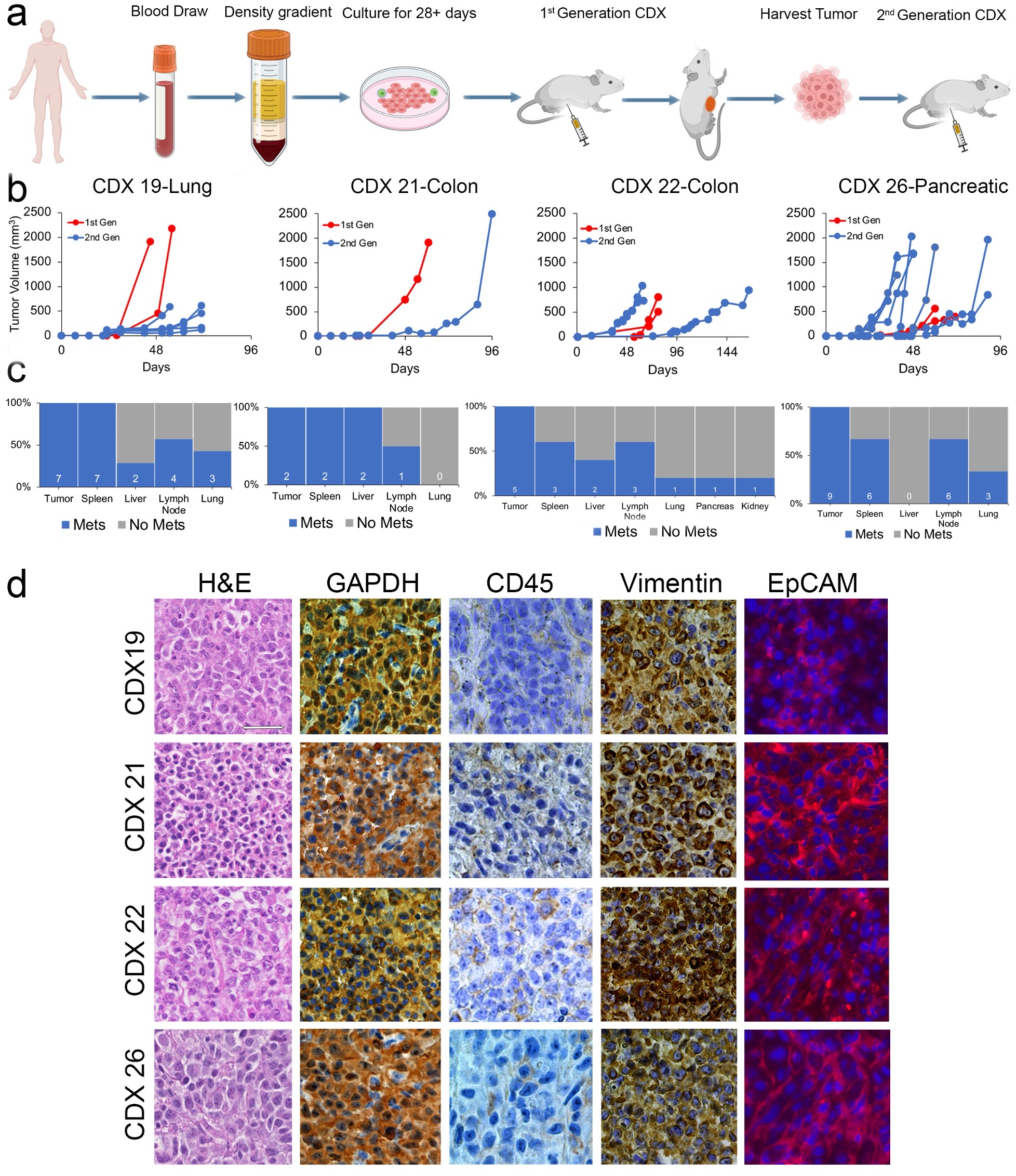
Four patient derived CDX models were established following a period of short-term culture. (a) Schema of CDX model formation created using Biorender. (b) Growth charts indicating the progression of CDX tumors. 1^st^ generation CDX models were established via the injection of cultured CTCs; 2^nd^ generation CDX models were established via the serial re-engraftment of tissue derived from 1^st^ generation CDX primary tumors. (c) Corresponding bar graphs indicating the number of CDX models with evidence of tumor in distant organs, separated by patient. (d) Representative hematoxylin & eosin (H&E, column 1), DAB chromogen staining of human specific GAPDH (column 2), common leukocyte marker CD45 (column 3), and vimentin (column 4), and immunofluorescence staining for EpCAM was performed on 1^st^ generation CDX primary tumors. Scale bar = 25 um.

Patient #19 was a 56-year-old male diagnosed with stage IV non-small cell lung adenocarcinoma with confirmed metastases to the vertebral bone. He was initially treated with 30Gy radiation to the vertebral lesion and systemic first-line combination chemotherapy (carboplatin and pemetrexed) and immunotherapy (pembrolizumab) with eventual progression of metastatic bony disease requiring vertebral surgery and femur surgery and radiation. A liquid biopsy for CTC culture was collected before initiation of second-line systemic therapy with docetaxel and ramucirumab. The patient ultimately developed numerous progressive hepatic and osseous metastases after 3 cycles of therapy, endured multiple hospitalizations for cord compression and cholangitis, and died 9 months after the initial diagnosis

Patient #21 was a 69-year-old male diagnosed initially with stage II sigmoid colon adenocarcinoma treated with curative-intent surgical resection and adjuvant chemotherapy. After two years, he was diagnosed with stage IV disease with oligometastatic recurrence in the soft tissue between the bladder and rectum that was HER2-amplified, microsatellite stable, and RAS and BRAF wildtype. He progressed locally on 2 lines of systemic therapy, underwent oligometastectomy with partial cystectomy and en bloc colon resection, but eventually had systemic progression in the lungs. He failed to respond to third-line HER-2 directed therapy. Liquid biopsy was collected for CTC culture during fourth-line systemic treatment on which he ultimately had progression of disease.

Patient #22 was a female with an 18-year history of ulcerative colitis who was diagnosed at age 31 with stage IV sigmoid colon cancer characterized as RAS and BRAF wildtype, HER-2 non-amplified, and microsatellite stable with metastases to the liver and pelvis. A liquid biopsy was collected for this study prior to initiation of first-line systemic treatment with FOLFOX and panitimumab. She ultimately underwent surgical resection of pelvic disease and a total abdominal colectomy after disease response to systemic therapy but ongoing ulcerative colitis symptoms and subsequently achieved stable disease status on maintenance systemic therapy for 12 months. Eventually, she had progression of disease in the liver, abdominal lymph nodes, peritoneum, and right atrium treated with various local surgeries and systemic therapy. She died after progression on fifth-line systemic therapy about 2 years after initial cancer diagnosis.

Patient #26 was a female diagnosed at age 82 with stage IV moderately-differentiated pancreas adenocarcinoma with metastases to the liver, lymph nodes, and lung. A liquid biopsy was collected for CTC culture prior to initiation of first-line, palliative, systemic treatment with gemcitabine and nab-paclitaxel. She only received one dose of systemic therapy after which her course was complicated by transaminitis and hyperbilirubinemia due to progressive liver disease resulting in biliary obstruction that was unamenable to local intervention. She was recommended best supportive care, pursued home hospice, and ultimately died 3 months after her initial diagnosis.

To establish CDX models, CTCs separated from whole peripheral blood were first cultured for 28 days as described in **Fig. 2a**. CTCs expanded in culture (5,000 to 25,000 cells) were injected subcutaneously into the flanks of immunodeficient mice. These mice are referred to in this report as “1^st^ generation CDXs.” All 1^st^ generation CDXs developed palpable primary tumors within 32 ± 6 days post-injection (n = 7 mice) and were euthanized by day 78 (mean 65 ± 30 days) (**Fig. 2b**). Tumor tissues from 1^st^ generation CDXs were divided and reimplanted into another mouse of the same immunodeficient strain to generate the “2^nd^ generation CDXs.” 2^nd^ generation recipient mice developed palpable primary tumors at the site of injection within 31 ± 10 days (n = 16 mice) and were euthanized by day 165 post-transplantation (mean 75 ± 29 days) (**Fig. 2b**). Among all 1^st^ and 2^nd^ generation CDX models, macro-metastatic nodules visible to the naked eye were observed in 21 of the 23 models on autopsy, involving an average of two distant organs per mouse (**Fig. 2c**). Overall, the spleen was the most common site of metastasis (19/21 90.5%), followed by the axillary lymph nodes (14/21, 66.7%), lungs (7/21, 33.3%), liver (6/21, 28.6%), kidneys (1/21, 4.8%) and pancreas (1/21, 4.8%) (**Fig. 2c**). Hematoxylin-eosin staining of tumors and macro-metastases indicated poorly differentiated carcinoma cells (**Fig. 2d**). IHC staining using human-specific antibodies targeting GAPDH, VIM as well as immunofluorescent (IF) staining for EpCAM all demonstrated positivity in both CDX tumors and metastatic nodules. No detectable staining of human CD45 was observed in CDX tissues (**Fig. 2d**).

Recently, cases of spontaneous lymphomagenesis in immunocompromised mice following xenograft of human cells have been reported^36^. Therefore, we sought to dispel this possibility in the CDX mouse models. Following isolation of RNA from CDX tissues and RNA sequencing, we used the CIBERSORT software^37^, ProteinAtlas^38^, and reactome pathway analysis to identify proportions of human immune cells, comparing RNA obtained from the whole blood of the patient and the corresponding tumor tissue of the CDX model. CIBERSORT showed that CDX tissues were significantly depleted for not only CD4^+^ naïve and CD8^+^ T-cells, but also B-cells, natural killer (NK) cells, and monocytes (**Fig. S2a**). Immune-cell specific signatures derived from the ProteinAtlas^38^ also indicated a decrease in expression of genes specific for B and T cells, NK cells, dendritic cells, macrophages, plasma cells, and granulocytes (**Fig. S2b**). Finally, reactome pathway analysis demonstrated depletion of human genes associated with the innate immune system in cultured CTCs and CDX tissues relative to samples obtained from healthy donors (**Fig. S2c**). Taken together these findings rule out the presence of murine spontaneous lymphomagenesis.

### CTCs and CDXs express transcriptomic heterogeneity

To evaluate transcriptomic similarities or differences between healthy donor whole blood (HD), matched patient-specific whole blood (WBM), cultured CTCs (TC), and CDX models, bulk RNA-sequencing was performed on all samples, and differential gene analysis carried out (**Fig. 3a**). For analysis, HD and WBM samples were pooled together and compared against TC and CDX tissues. Compared to HD and WBM, 848 genes were significantly (p-value < 0.05) upregulated and 21 genes were significantly downregulated in TC and CDX models. When TC were compared with WBM, 1828 and 1165 genes were significantly up- or down-regulated, respectively (**Table S1**). Similarly, 3129 and 3419 genes were significantly up- or down-regulated, respectively in CDX tissues compared to WBM (**Table S1**). Finally, 1836 and 2731 genes were significantly up- or down-regulated in CDX tissues compared to TC (**Table S1**). These data summarize pooled data for all four patients; patient-specific comparisons are further provided in **Table S1**.

**Figure 3:**
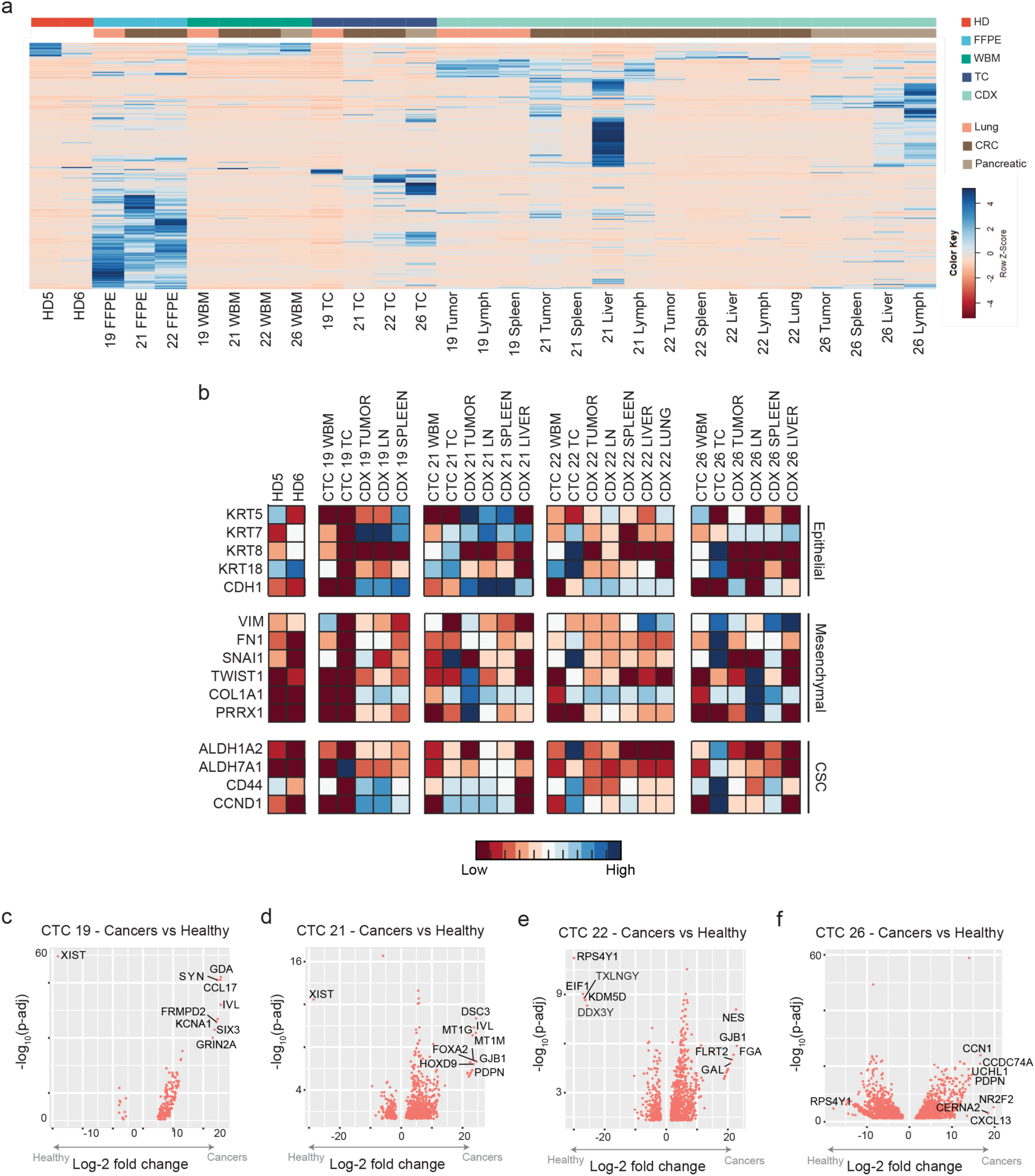
Bulk RNA-sequencing on cultured CTCs and CDX tissues show transcriptomic heterogeneity. (a) A clustergram depicting mRNA expression in the various samples evaluated. (b) A panel of epithelial (KRT5, KRT7, KRT8, CDH1, EpCAM), mesenchymal (VIM, FN1, SNAI1, TWIST1, COL1A1, PRRX1), and cancer stem cell (ALDH1A2, ALDH7A1, CD44, CCND1) was extracted. In general, CTC-derived samples demonstrate increased expression relative to healthy donor and whole blood samples. (c-f) Volcano plots for all significantly differentially expressed genes between patient-specific cancer and healthy samples are plotted. A select few of the most statistically significant differentially expressed genes are labeled for each patient.

Previous studies including ours have identified activation of the epithelial-mesenchymal transition program as a critical event in the transition of a cancer cell from primary tumor to metastasis^31^ include ours. We therefore evaluated the expression of a panel of epithelial (KRT5, 7, 8, 18, CDH1), mesenchymal (VIM, FN1, SNAI1, TWIST1, COL1A1, PRRX1), and cancer stem cell (ALDH1A2, ALDH7A1, CD44, CCND1) markers in all samples. We observed increased expression of markers associated with metastases in CTC-derived samples (TCs and CDXs) compared to HD and WBM (**Fig. 3b**).

Lastly, patient-specific differential gene analysis was performed comparing the TCs and CDX tissues with HD and WBM samples (**Fig. 3c-f**). Notable genes that were upregulated in patient-derived CTC models include GDA, FOXA2, GJB1, and NR2F2 (**Table S2**). On the other hand, XIST and RPS4Y1 were the most downregulated genes in each respective patient sample (**Table S2**).

### Five genes are associated with high-risk CTCs

Recent studies have indicated that CTCs exhibit considerable transcriptional heterogeneity^13^. A robust gene biomarker across multiple cancer types for CTCs has yet to be identified^39, 40^. Therefore, we interrogated our data to identify genes whose expression were at least eight-fold higher in TCs or CDX tissues compared to the matched WBM. Based on this criterion, 199 (**Fig. 4a, Table S3**) and 35 (**Fig. 4b, Table S4**) genes were identified from cultured CTCs or CDX tissues, respectively. Of these, we identified eight genes (*IGSF3, EMX1, TFPI2, CCL22, BCAR1, RRAD, CCL1, COL1A1*) (**Fig. 4c**) that were upregulated in all CTC cultures and CDX tissues. Kaplan-Meier survival analyses using TCGA pan-cancer atlas clinical data and gene expression data of these eight genes further narrowed the signature to a subset of five genes (*BCAR1, COL1A1, IGSF3, RRAD, and TFPI2*) whose increased expression was significantly associated with poorer disease--free survival (Fig 4d and **Table S5**).

**Figure 4:**
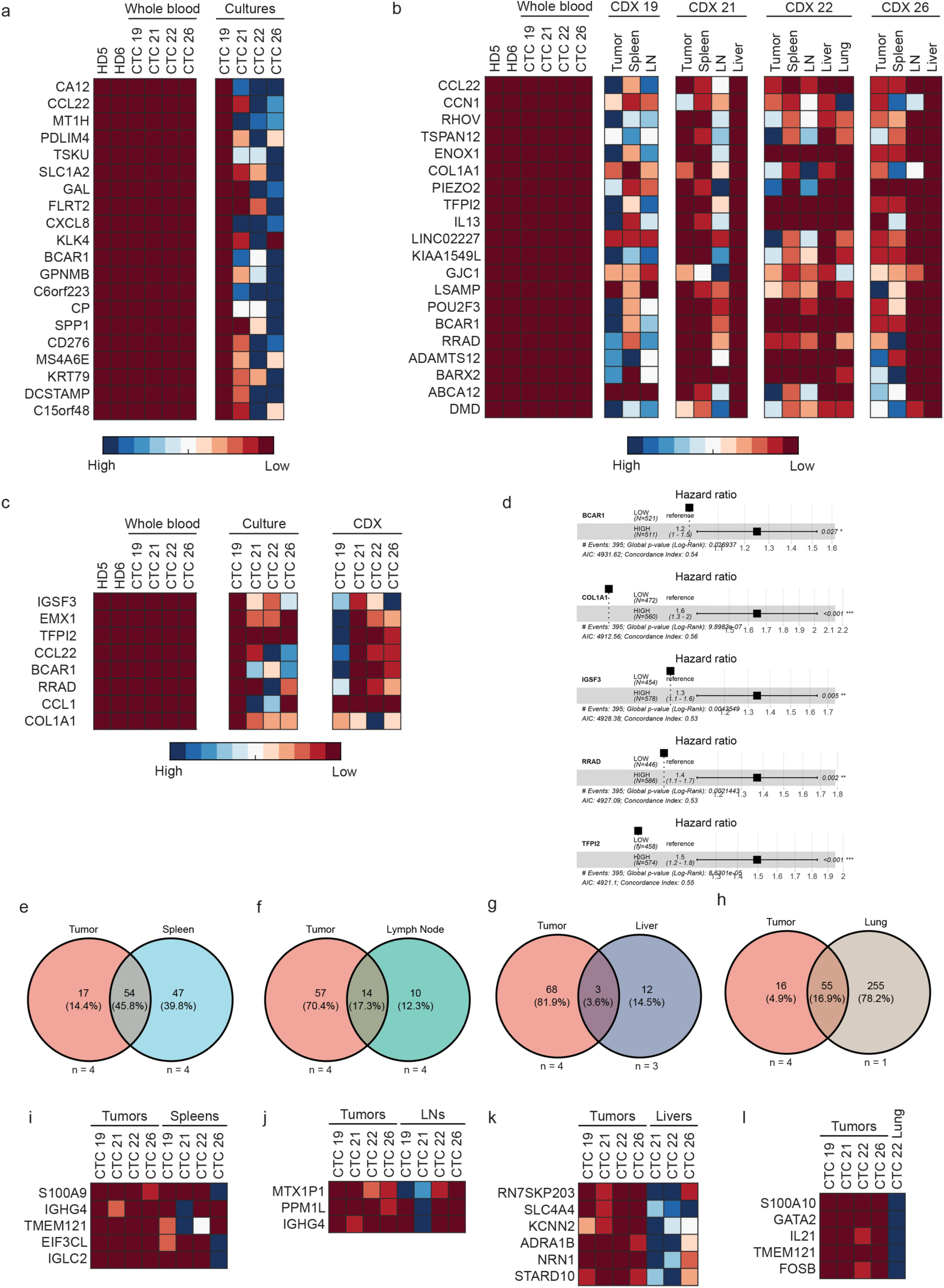
A prospective CTC-specific signature was identified. Differential gene analysis was used to determine a 35 gene signature (top 20 most differentially expressed depicted here) (a) and a 199 gene signature (b) that could reliably identify any CTC cultures or CDX tissues, respectively. (c) A final gene signature composed of eight genes (IGSF3, EMX1, TFPI2, CCL22, BCAR1, RRAD, CCL1, COL1A1) that were routinely over-expressed in any CTC-derived sample was identified. (d) COX proportional hazards ratios were calculated based on gene expression of the 8 CTC-signature genes and overall survival within the relevant TCGA pan-cancer atlas databases. Higher expression of five of the eight genes was associated with a poorer overall survival. (e-h) Differential gene analysis was performed on an CDX organ-specific level relative to a common control of whole blood samples. Venn-diagrams depict the amount of overlap in genes that were overexpressed in tumors and genes that were overexpressed in the various respective distant organs. Organ-specific metastatic signatures were then identified by comparing expression of genes in CDX tumor samples and respective distant organs. Ultimately, (i) a four-gene spleen metastatic signature, (j) two-gene lymph node metastatic signature, (k) six-gene liver metastatic signature, and (l) five-gene lung metastatic signature were identified.

In addition to a gene signature for CTCs, we also sought to determine if our CDX models could be used to identify prospective organ-specific metastatic signatures. For instance, transcriptomic signatures for lung and brain-specific breast cancer have been identified through the evaluation of matched xenografted tumors and resultant metastatic leisons^41, 42^. Gene expression was compared between CDX primary tumors and their respective lymph node, spleen, lung, and liver metastases. Across organs, 47, 10, 12, and 255 genes were uniquely expressed in the CDX spleen, lymph node, liver, and lung that were not overexpressed in the primary CDX tumor (**Fig. 4e-h**). To further narrow the prospective organ-specific signatures, we set a threshold expression of greater than eight-fold in the metastases compared to the CDX primary tumor. In all, 4, 2, 6, and 5 genes met this criterion in spleen, lymph node, liver, and lung metastases respectively resulted in the final identification of four prospective organ-specific signatures (**Fig. 4i-l**).

### NF-kB signaling as a biomarker for CTCs

Another method of evaluating transcriptomics involves the differential analysis of gene sets and signaling pathways through the Kyoto Encyclopedia of Genes and Genomes (KEGG)^43^ and Gene Set Enrichment Analysis (GSEA)^44^. Patient-specific analysis comparing signaling pathways enriched in cultured CTCs or CDX tissues compared to WBM was performed using KEGG. Cultured CTCs and CDXs from the four patients showed enrichment of the following signaling pathways among others: Patient #19, mTOR1, NOD-like receptor and NF-kB (**Fig. 5a**); Patient #21, TNF, PI3K-AKT, IL-17, and NF-kB (**Fig. 5b**); Patient #22, Chemokine, p53, NF-kB, IL-17, and TLR (**Fig. 5c**); Patient #26, PI3K-Akt, TNF, and NF-kB (**Fig. 5d**).

**Figure 5:**
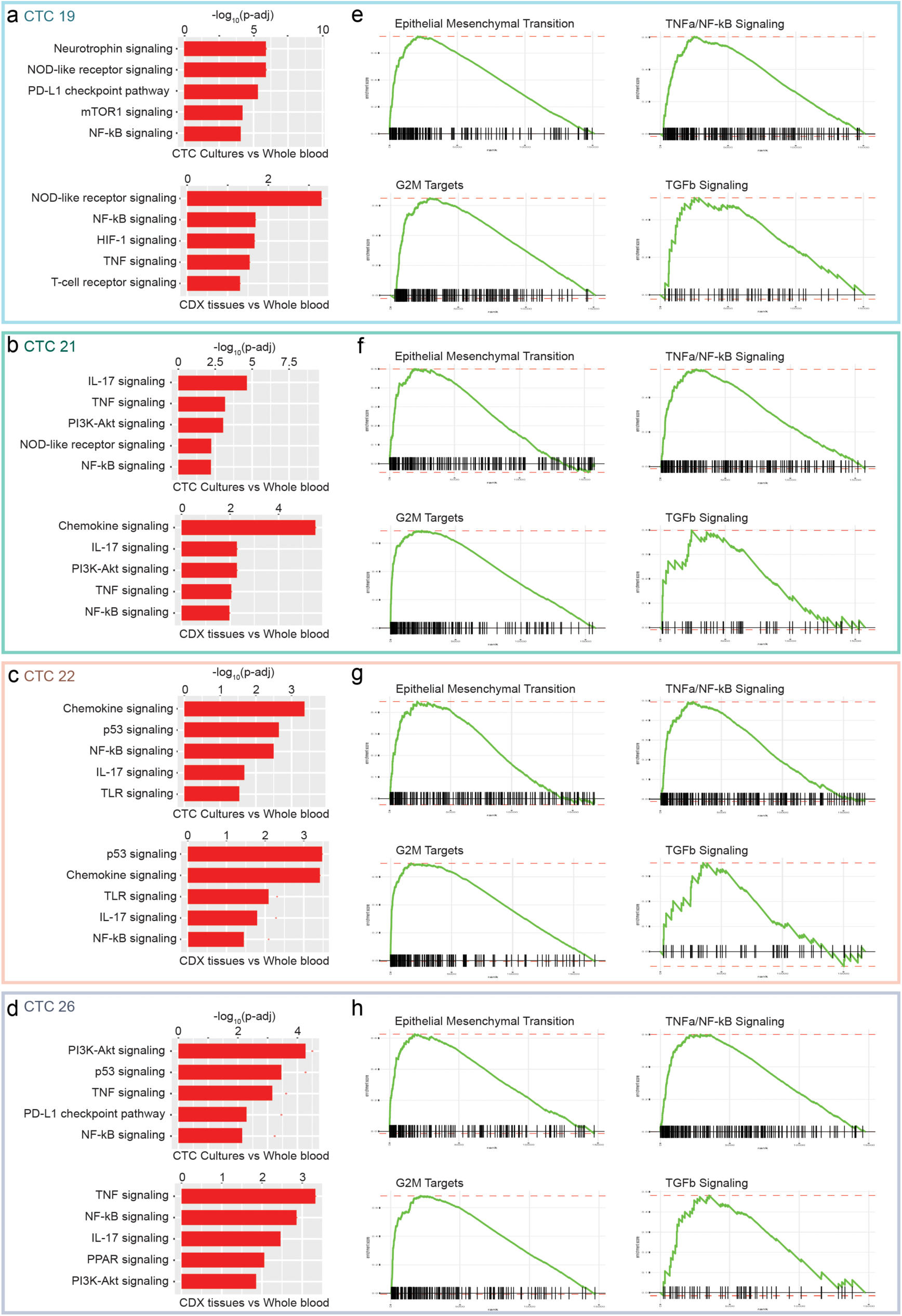
KEGG and GSEA pathway analysis on CTC-derived samples indicate an enrichment of NF-kB signaling. Waterfall plots of the top 5 pathways enriched via KEGG analysis in patient-specific CTC cultures or CDX tissues against whole blood samples for (a) patient #19, (b) patient #21, (c) patient #22, and (d) patient #26 are depicted. GSEA enrichment plots depicting enrichment of the hallmarks genesets EMT, TNFα/NF-kB signaling, G2M targets, and TGF-β signaling are shown for samples from patient #19 (e), patient #21 (f), patient #22 (g), and patient #26 (h).

Notably, CTCs are composed of a highly heterogeneous population of cells^45, 46^. While pathway analysis of cultured CTCs and CDXs revealed enrichment of shared signaling pathways, it was not surprising that differences in enriched signaling pathways were also observed, which can likely be attributed to the microenvironment that the CTCs are present in. For instance, PD-L1 and mTOR1 signaling were enriched in cultured CTCs from patient #19, but not the derivative CDX tissues (**Fig. 5a**). Conversely, HIF-1 and TNF signaling were enriched in CDX tissues derived from patient #19, but not the source cultured CTCs. Similarly, other individually enriched signaling pathways were observed in either CDX tissues (e.g., chemokine signaling in CDX tissue from patient #21) or cultured CTCs (PDL-1 signaling and p53 signaling in CTC from patient #26) only (**Fig. 5b-d**).

GSEA pathway analysis offers a more focused approach to cancer, evaluating 50 sets of genes associated with specific biological states or processes with minimal redundancy^47^. As described above, we compared the gene expression from TCs and CDX tissues with WBM from the same patient. The geneset that defines the epithelial-mesenchymal transition was significantly enriched in all four patients. Similarly, TNFα signaling via NF-kB was also enriched in models derived from all four patients. Both EMT and NF-kB signaling are routinely identified in studies evaluating cancer progression and metastasis^48^. Other genesets that were routinely enriched in CTC-derived samples were G2M targets and TGF-β signaling, among other genesets (**Fig. 5e-h** and **Fig. S3**). These results are consistent with the findings from the KEGG analyses.

Furthermore, a functional network was generated through input of the 199 gene prospective TC signature from **Table S3** into String-DB, using the whole human genome as background. Terms found to be enriched across the CTC gene signature were selected, and member proteins overlapping those processes and the CTC signature were plotted. Nodes represent all proteins produced by a single, protein-coding gene locus, either by associated gene ontology (GO), KEGG pathway, or WikiPathway (WP) terms. All enriched terms showed a FDR < 0.05 (**Fig. S4**). Notably, core nodes within this signature involved the PI3K-Akt, MAPK, and NF-kB signaling pathways. Together, these data suggest NFκB signaling as a biomarker for metastasis.

### CDXs can be queried for metastasis-driving mutations

In addition to transcriptomic heterogeneity, CTCs are also known to acquire *de novo* mutations separate from their respective primary tumors^49, 50^. In some cases, these *de novo* mutations have been identified as potential drivers of resistance to therapeutics^24, 51^. Additionally, multiple *de novo* CTC-exclusive mutations have been identified in genes that are associated with human cancers, namely*, KRAS, BRAF*, and *PIK3C*^52, 53^.

To evaluate the prevalence of mutational concordance/discordance in our samples, we performed whole exome sequencing (WES). WES analysis of HD, patient-specific FFPE, TCs, CDX primary tumors, and corresponding distant metastatic CDX lesions was performed and compared against pathology reports from primary tumor biopsies obtained from patients where available (patients #21, 22, 26). In patient #21, of the 7 single nucleotide polymorphisms (SNPs) reported in the primary tumor, 5 were identified in the TC and CDX primary tumor (**Table 1**). In patient #22, a SNP in the cancer tumor suppressor gene *TP53* reported in the patient’s primary tumor was identified in the corresponding FFPE and CTC culture, but not the CDX primary tumor or distant metastases (**Table 1**). Finally, in patient #26, a SNP in *SF3B1* reported with a 11% variant frequency in the patient’s primary tumor biopsy was identified in the matched CTC, CDX and CDX metastases (**Table 1**).

**Table 1.**
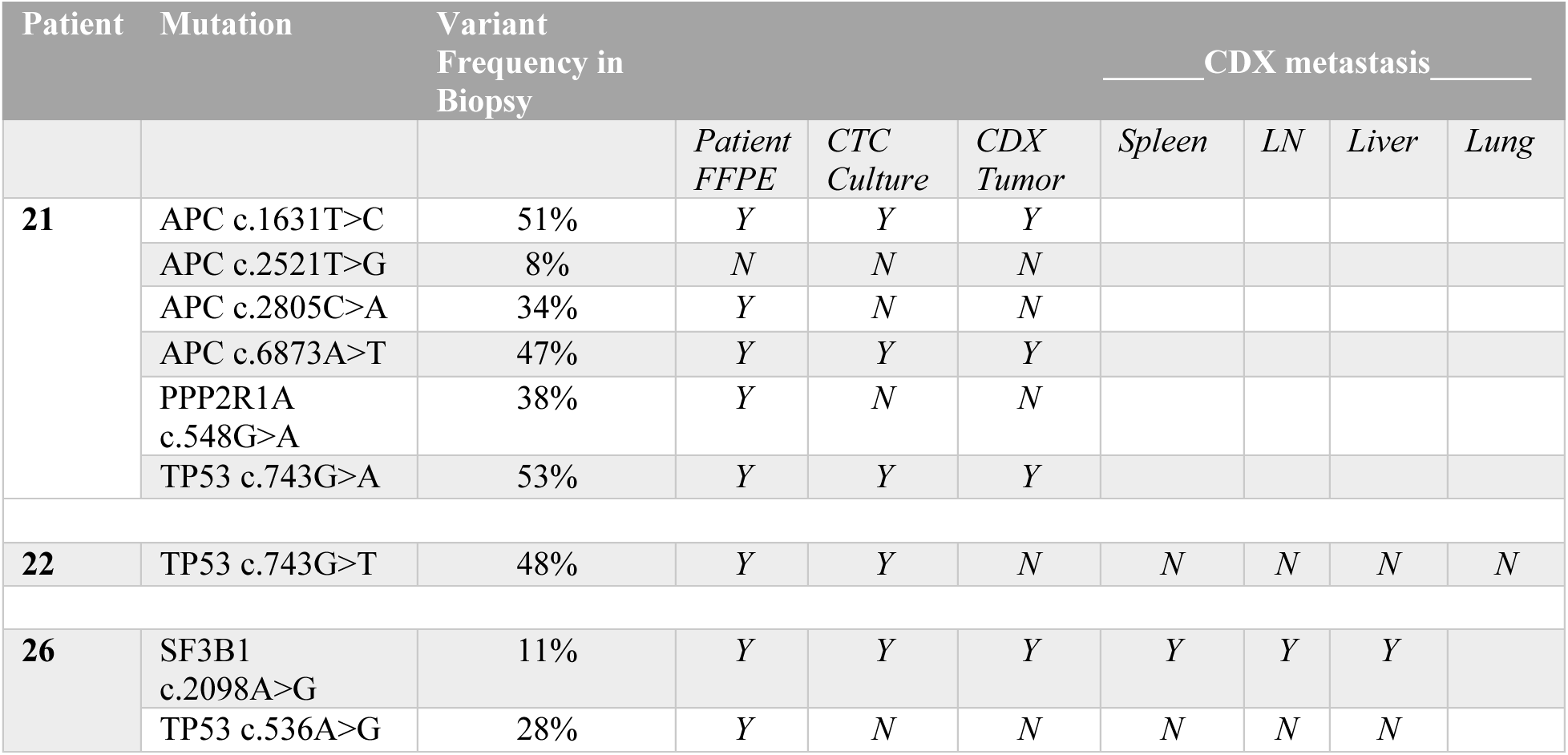
Single nucleotide polymorphisms were compared between clinical samples of patient primary tumor biopsies and corresponding FFPE, CTC cultures, CDX tumors, and CDX metastatic organs.

Next, a mutant allele fraction (MAF) was calculated to determine the presence of true SNPs within our samples (**Fig. 6a**). Unfortunately, DNA sequencing of patient’s FFPE tissue samples resulted in poor quality reads across a large majority of the genome, representing difficulties in whole-exome sequencing of patient tumors (**Fig. 6a**). Despite these limitations, across all four samples, ≥ 80% of all FFPE identified mutations were recovered in corresponding TC and CDX models, indicating strong genetic fidelity across models (**Fig. 6a**). On the other hand, some variability in mutations across samples did exist (**Fig. 6b-e)**. These findings ultimately confirm genetic fidelity across the two distinct model systems (TC and CDX) as well as demonstrates mutational heterogeneity consistent with the known behaviors of CTCs.

**Figure 6:**
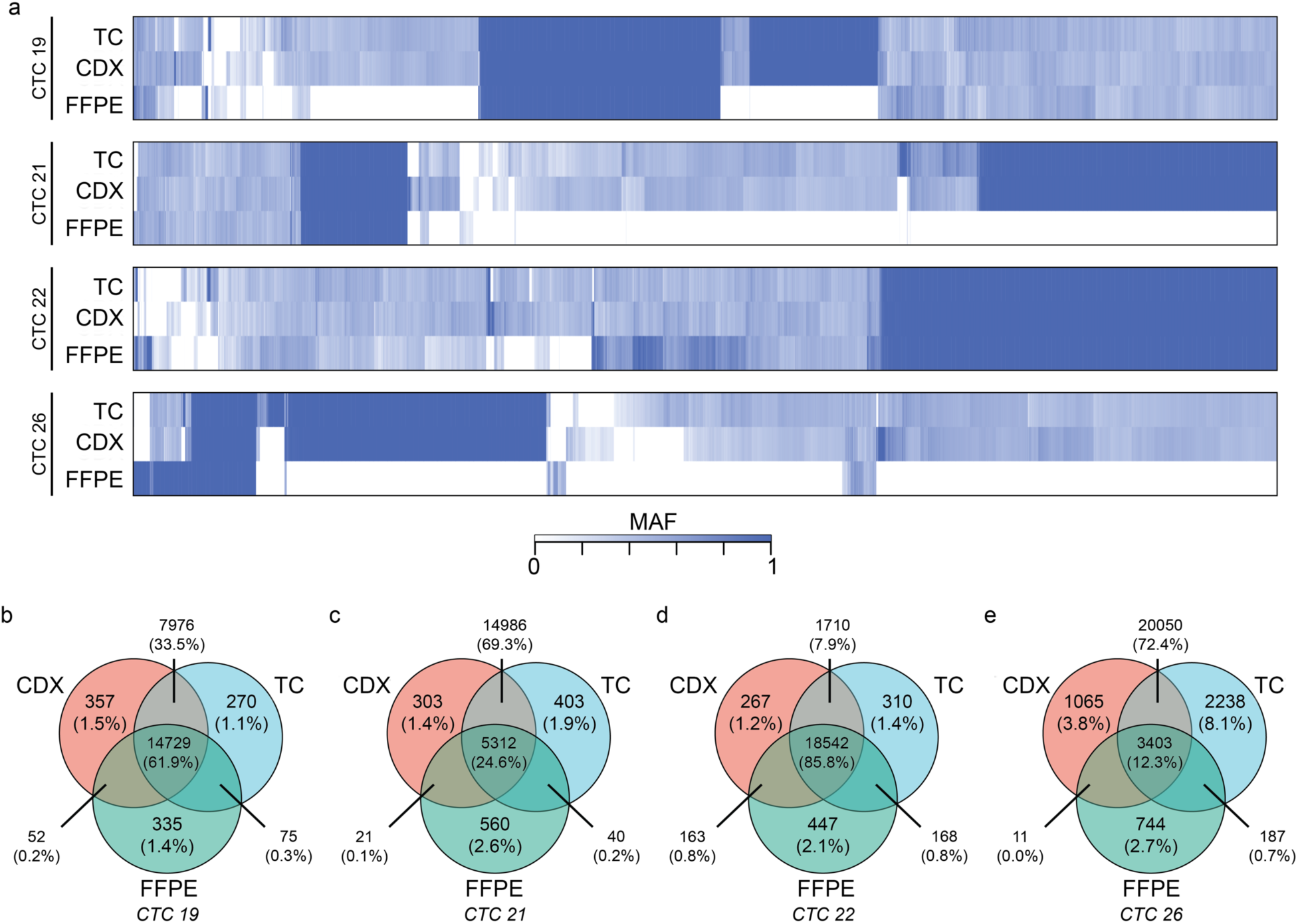
Whole exome sequencing revealed concordance and discordance between patient’s tumor and corresponding CTC models (TC and CDX). (a) Heatmaps depicting the mutant allele fraction (MAF) of all mapped mutations across patient-matched CTC cultures (“TC”), CDX primary tumors (“Tumor”), and clinical formalin-fixed paraffin embedded (“FFPE”) tissue. More mutations were recovered from CTC and CDX tissues than FFPE, on average. A MAF indicates homozygous mutation, while 0.5 indicates heterozygous mutation. (b-e) Mutations were then mapped in a Venn diagram to determine the number of overlapping and sample-exclusive mutations for each patient.

Next, we hypothesized that *de novo* oncogenic mutations identified in cultured CTCs or CDX primary tumors could signal a role in metastasis. The OncoKB database, curated from world-wide sources, provides detailed information regarding specific alterations in 682 cancer genes^54^. In 2021, the FDA recognized a portion of the OncoKB as a source of valid scientific evidence for Level 2 (clinical significance) and Level 3 (potential clinical significance) biomarkers. This means that submissions to the FDA can use these data to support the clinical validity of tumor profiling tests. In patient #19, fifteen oncogenic mutations were CDX tumor-exclusive, affecting genes such as *CDK8, COP1, and MAP2K2* (**Table S6**). In patient #21, mutated genes in cultured CTCs included *AKT1, SERPINB3, EGFR, and NOTCH4*, while *COP1* was once again mutated in CDX tumors (**Table S7**). In CDX tumors derived from patient #22, *COP1* and *GNAS* were mutated, while *NUF2* was the only oncogenic mutation identified in CTC cultures (**Table S8**). Multiple *de novo* mutations in *COP1, CDK8, and AKT2* were observed in CDX primary tumors derived from patient #26, while mutations in *SERPINB3, MGAM*, and *HDAC7* were observed in corresponding patient-matched cultures (**Table S9**). In summary, mutation in *COP1* was identified in all four CDXs from diverse tumor types suggesting *COP1* as a potential driver of metastasis.

Mutations in specific genes could also be responsible for organ-specific patterns of metastasis^48^. We therefore sought to determine whether a consistent pattern of mutations driving metastasis could be identified based on specific organs (**Fig. 7a**). No WES analysis was conducted on distant organs from CDX models derived from patient 21 due to poor quality sequencing. When considering all CDX primary tumors, 60.4% of all identified mutations were unique to a single patient-derived primary tumor, while approximately 5.6% of all mutations occurred in all four primary tumors sequenced (**Fig. 7b**).

**Figure 7:**
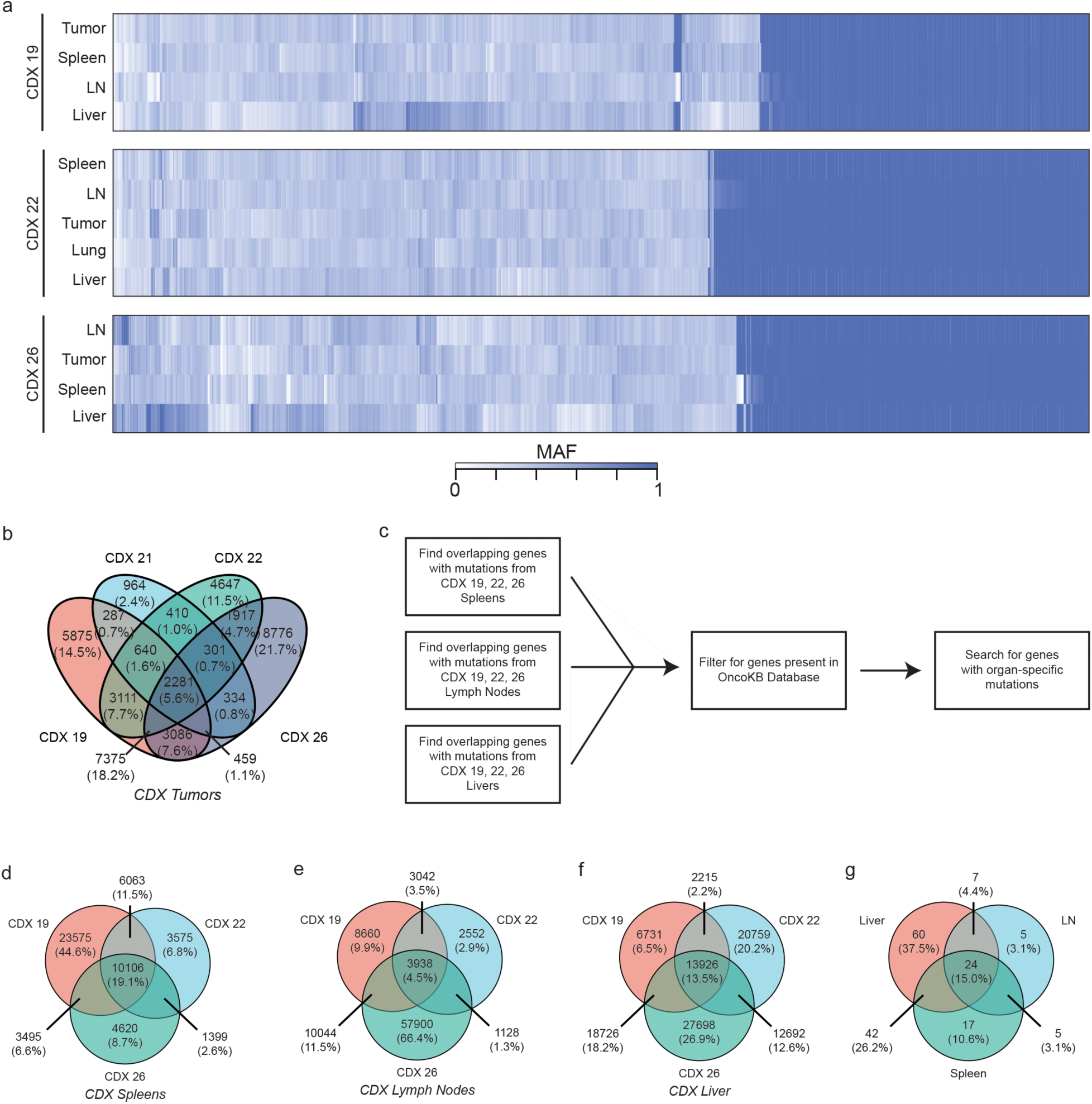
Distant metastases in CDX models were queried to identify organ-specific mutations. (a) Heatmaps based on patient specific CDX models indicate that a large proportion of mutations were consistent across CDX primary tumors and distant metastases. (b) Mutations across all CDX tumors were mapped to identify the percentage of mutations that were consistent across all CDX tumors. (c) A schema for identifying organ-specific mutations is shown. Mutations were mapped to identify distribution of unique mutations across all CDX (d) spleens, (e) lymph nodes, and (f) livers which had evidence of metastases. (g) A Venn diagram evaluating for unique genes with mutations depending on organ evaluated is shown.

To identify potential organ-specific metastatic mutations/genes, we developed an analytical workflow that is depicted in **Fig. 7c**. Across all unique mutations in CDX spleens, 19.1% of mutations were shared across all three spleen samples queried (**Fig. 7d**). Similarly, 4.5% and 13.5% of all unique mutations were shared across all three CDX lymph node or liver samples, respectively (**Fig. 7e-f**). After filtering for only those genes that were included within the OncoKB database, we identified 60 unique genes, including *MTOR, NOTCH2, JAK1, JUN* and *HIF1A*, that harbored mutations only within the liver samples and were not present within lymph nodes or spleen samples (**Fig. 7g** and **Table S10**). On the other hand, 5 genes (*KDR, RUNX1T1, RPS6KB2 NOTCH3, EPOR*) exhibited mutations exclusively in CDX lymph node tissues. Finally, 17 genes exhibited mutational exclusivity in CDX spleens, including *YAP1, HDAC7, MAPK3*, and *SOX9* (**Fig. 7g** and **Table S10**).

## DISCUSSION

Circulating tumor cells could be key to understanding the metastatic cascade and developing directed therapies against metastasis. This study introduces a novel platform for isolating and culturing CTCs *ex vivo* that significantly improves upon previous attempts. Leveraging an unbiased density-dependent isolation process that is independent of selection by antibodies or microfluidic devices, we have established viable CTC cultures from 23/25 total samples spanning lung, colon, and pancreatic adenocarcinomas (**Fig. 1**), confirming the high yield previously described by us with breast cancer clinical samples^27^. In addition, the expanded CTC cultures were successfully used here to generate robust CDX models with extensive macro metastases.

Previous attempts at culturing CTCs *ex vivo* have been hampered by the requirement of high numbers of CTCs isolated directly from whole blood^11, 17, 18^. Historically, antibody-dependent isolation methods have been used to concentrate CTC^15, 55^. However, in most patients CTCs are ultra-rare (< 10 cells/mL)^15^. For instance, isolation and culture of CTCs from patients diagnosed with gastroesophageal cancers using the commercially available RosetteSep® could only establish CTC cultures from 2 out of 41 (a 4.9% success rate) patients, both of whom had at minimum 109 CTCs/7.5 mL of blood^22^. Such a high CTC count is rare in even the most metastatic cancer patients. In addition, most previous reports of CTC cultures have been targeted towards the culture of CTCs derived from a single cancer type^11^. In contrast to previous studies, in this report we demonstrate successful expansion of cultured CTCs from regular peripheral blood draw in patients diagnosed with one of three distinct tumor types (colon, lung, and pancreatic adenocarcinomas) using a single generalized culture technique (**Fig. 2a**). This culture system enables the harvest of CTCs along with CTC-supporting cells that may be present within whole blood^27^.

Similarly, in previous attempts to establish CDX models, only those mice infused with samples from patients with very high CTC counts (≥ 400) successfully established CDXs^18, 27, 33–35^. Given the rarity of CTCs, seeking patients with high numbers of CTCs in the blood is not a viable strategy for the routine initiation of CDX models. Our ability to successfully expand CTCs in culture with high success rate (92%) offered a plausible strategy to routinely initiate CDX models using ≧5,000 CTCs/injection opening the use of this technology to initiate both CTC short-term cultures and long-term CDXs in a large cohort of patients spanning across multiple cancer types. In the proof-of-concept described here, we used CTCs from 4 patients with diverse cancers that were first expanded through in short-term cultures (14-28 days) to attain numbers suitable for the establishment of patient-derived CDXs (**Fig. 2**). We not only successfully established CDX models using cultured CTCs from all four patients, but also demonstrated replicable patterns of metastasis through multiple generations. Of the few CDX models previously described, many fail to demonstrate metastasis, only generating a tumor at the site of injection^25, 34, 35^ with high latency period of at least 6 months. The four CDX models generated here all exhibited macro-metastatic nodules in distant organs within 2-3 months. Our models thus recapitulate the physiological function of CTCs as a metastasis-initiating population.

While our primary purpose in expanding CTCs *ex vivo* was to enhance the numbers of CTCs available for initiating CDX models, our data suggest that it is also plausible that the short-term culture selected for a highly aggressive and adaptable CTC population that was particularly suited to survival in adverse conditions such as *ex vivo* culture. For instance, cultured CTCs were enriched in expression for not only epithelial markers, but also mesenchymal and cancer stem cell markers (**Fig. 2-3 and Fig. S1**). This expression of EMT plasticity in CTCs is of particular interest, as it has been demonstrated that mesenchymal-expressing CTCs are more likely to cluster and form metastatic lesions *in vivo*^12^. Furthermore, previous single-cell RNA sequencing of isolated CTCs demonstrated that CTCs are a highly dynamic and heterogeneous population^12^. Genomic heterogeneity seen among CDX tumor and its corresponding metastases suggest that at least some level of heterogeneity waa maintained in the cultured CTCs and CDX models (**Fig. 6-7 and Tables S6-S10**) Further studies plan to evaluate how culturing and expanding CTCs affects the composition of the heterogenic population at the single cell level.

We leveraged the patient-derived CTC cultures and CDX models to determine enrichment of specific molecular pathways and gene signatures in cultured CTCs and CDXs derived from the in a pan-cancer cohort. In a previous study, we identified enrichment of TNFα signaling via NF-kB in CTC cultures derived from breast cancers^27^. In this study, TNFα signaling via NF-kB was once again a commonly enriched pathway in cultured CTCs and their corresponding CDX models derived from all 4 patients (**Fig. 5**). Thus, the repeated identification of TNFα suggests that it is a critical and conserved pan-cancer pathway involved in metastasis. Other reports have also identified TNFα signaling in metastasis in a variety of cancers, identifying perturbations associated with enhanced metastatic potential^56^. In the clinical setting, anti-TNFα inhibitors are now entering clinical trials to treat cancers^57^. The indirect targeting of TNFα signaling through NF-kB inhibitors has also been more widely studied in cancers, as NF-kB is frequently constitutively activated in most human cancers^58^. In the non-canonical pathway, NF-kB activates a signaling cascade that works through the TNF receptor to introduce a set of cytokines, eventually resulting in activation of inflammation^58^. Several pharmacological strategies have been employed to target NF-kB in cancers, including inhibition of upstream IKK phosphorylation, immunomodulatory agents such as thalidomide, or proteasome inhibitors such as bortezomib^58^. Existing evidence of the importance of TNFα in human cancers is therefore supplemented with the novel findings in the current study. Taken together, these studies strongly suggest that direct or indirect targeting of TNFα signaling could be a potentially potent strategy for developing metastasis-specific therapeutic agents. The routine establishment of individual patient-derived CTC cultures and matched CDX models will provide a key biological resource for screening molecules that target TNFα as specific anti-metastatic agents.

In this study, differential gene expression analysis revealed a prospective CTC signature consisting of five genes (*BCAR1, COL1A1, IGSF3, RRAD, and TFPI2*) whose expression (a) was robustly enriched across all CTC-derived samples (**Fig. 4c**), and (b) was associated with poorer disease-free survival using pan cancer TCGA dataset (**Fig. 4d**). Notably, all five genes that constitute the metastatic signature have been previously individually linked to metastatic phenomena.

BCAR1 has previously been identified as a novel binding partner of mutant TP53, ultimately promoting cancer cell invasion^59^. Collagen type I alpha 1 (*COL1A1*) is perhaps the most strongly associated with the development of metastases in cancer as a critical element of the tumor microenvironment^60, 61^. IGSF3 acts as a promoter of hepatocellular carcinoma progression and metastasis through activation of the NF-kB signaling pathway^62^. Knockdown of Ras-related associated with diabetes (RRAD), a GTPase commonly implicated in metabolic function and hepatocellular carcinomas, suppressed cancer cell invasion, proliferation, and EMT^63, 64^. Finally, tissue factor pathway inhibitor 2 (TFPI2) has been shown to promote cancer metastasis through increased perivascular migration and ERK signaling^65, 66^.

Finally, we provide evidence of mutational discordance between patient primary tumors, FFPE, CTC cultures, and CDX tissues, demonstrating the *de novo* acquisition of mutations in oncogenic genes occurring during metastasis (**Figs. 6-7**). According to the classical viewpoint of cancer as a clonal disease, CTCs can develop an enhanced ability to metastasize due to the acquisition of novel genetic mutations^67^. While a large proportion of mutations were commonly shared between FFPE, CTC cultures, and CDX tissues, a focus on *de novo* mutations in cultured CTCs and CDX models could provide insight into prospective drivers of metastasis. For instance, mutations in constitutive photomorphogenesis protein 1 (*COP1*) were observed in all 4 CDX tumors evaluated (**Fig. 6 and Tables S6-S9**). In breast cancer cells, depletion of COP1 resulted in stabilization of the c-Jun protein and subsequent enhancement of cancer cell growth/migration and *in vivo* metastasis^68^. The prevalence of *COP1* across all four models could suggest a possible selective pressure for inactivating *COP1* mutations in metastasis. Interestingly, several other genes whose upregulation is commonly associated with increased metastatic potential also exhibited either culture or CDX model mutational exclusivity, including the serine-protease inhibitor *SERPINB3*, tyrosine kinase *EGFR*, and *AKT* isoforms 1 and 2 (**Fig. 6 and Table S6-9**). The latter is particularly interesting, as the various Akt isoforms are known to have varied effects on metastatic potential^69^. These findings may be partially supported by our own findings, which identified *AKT1* (an isoform generally associated with cancer cell growth and proliferation) mutations in cultured CTCs and *AKT2* (an isoform more commonly associated with metastasis) mutations in CDX tumors^69^. Finally, reports of CDK8^70^, EGFR^71^, MAP2K2^72^ and NOTCH signaling^73^ and *NUF2*^74^ mutations associated with enhanced metastasis are encouraging indications of the accuracy of the CTC-derived models established here.

This study provides evidence that CTCs from diverse cancers can be efficiently expanded *ex vivo*. More importantly, we demonstrate that such cultured CTCs can, in turn, be used to reliably, rapidly, and routinely establish CDX models. We have furthermore leveraged individual patient-derived CTCs and CDX tissue to genetically characterize CTCs. Scaling up these technologies would allow the establishment of sufficient numbers of CTCs and CDX models from individual cancers to identify biomarkers, and to screen existing and novel molecular entities for their capacity to target metastases. Patient derived CTCs and CDX models could also be used to design personalized therapeutic regimens for individual patients. Therefore, application of this novel platform to a broader cohort of patients could potentially lead to an improved understanding of the underlying mechanisms of metastasis as well as leverage CTCs as a key resource in developing a new generation of precision based personalized cancer treatments and vaccines for cancer patients including metastatic cancers.

## METHODS

### Patient Enrollment

Patients were recruited, consented, and enrolled at the Medstar Georgetown University Hospital Medical Oncology clinics in compliance with the Health Insurance Portability and Accountability Act (HIPAA) and Georgetown University Institutional Review Board (IRB) procedures (approval ID: MODCR00001156), managed through the Survey, Recruitment, and Biospecimen Collection shared resource (SRBSR) of the Lombardi Comprehensive Cancer Center. All patients provided written informed consent for the study. Inclusion criteria for patient enrollment included adult (> 18 years old), male or female patients with histologically confirmed metastatic disease and primary lung, colon and pancreatic cancers, with untreated or treated metastatic disease and with no limit of prior treatment lines. Exclusion criteria included any one or more of the following: (1) use of chemotherapy and/or antibody-based therapy less than a week before blood collection; (2) use of radiotherapy in the past 2 weeks unless there exist metastatic lesions beyond the radiated lesion. Healthy donors with no known health conditions at the time of consent were also enrolled. In all patients, 2 to 4 tubes of blood (∼7-8 mL/tube) were drawn. When applicable and available, blood was drawn from existing IV ports; if blood draws required skin puncture, an initial waste tube was disposed prior to processing. Deidentified pathology reports and clinical annotation regarding patient tumors and history were provided by the SRBSR.

### CTC Isolation from Whole Blood using FiColl-Paque

All patient peripheral blood samples were processed within 90 minutes of collection from patients. FiColl-Paque (Cytiva Life Sciences cat. no. 17-440-02) based separation of CTCs was performed as previously reported^27^. Briefly, samples were mixed with 1x Hank’s Balanced Salt Solution (HBSS) at 1:1 volume/volume ratio with whole blood at room temperature. 6 mL of the HBSS:blood sample was then split evenly into two 15 mL tubes containing 3 mL FiColl-Paque each, being careful not to mix samples with FiColl-Paque. Remaining HBSS:blood volume was split evenly into two 50 mL tubes containing 15 mL FiColl-Paque each. Samples were processed as described previously^27^. Upon completion of the final wash, cell pellets from 50 mL tubes were resuspended in 1 mL culture medium/tube. Cells were resuspended via gentle pipetting, and plated for short-term cultures. Cells resulted from 3 ml Ficoll-Paque tubes were used to extract DNA and RNA designated as whole blood match (WBM).

Blood processing for healthy donor blood proceeded in the same manner as isolation from enrolled patients. In healthy donors, two tubes of 7.5 mL blood each were drawn and processed via FiColl-Paque. Following all centrifugation and wash steps, cell pellets for healthy donors were reserved for RNA/DNA extraction only.

### CTC Cultures

Cells were resuspended in culture medium and plated for short-term cultures at 37°C for 14+ days as described previously^27^. Cultures were supplemented with fresh medium every 3 days and washed every 6 days with 1X PBS by centrifugation at 400x g for 4 minutes at 4°C. Every 6-10 days, cells were harvested via manual pipetting and resuspended in 1 mL of culture medium for automated trypan blue viability assays using the Thermo Scientific Invitrogen Countess II (AMQAX1000 cell counter). Phase microscopy images were taken using the EVOS FL Auto Imaging System (Thermo Fisher) at 20X zoom.

### CTC-derived Xenograft Models

All animal protocols and procedures for this study were approved by the Institutional Animal Care and Use Committee (IACUC) at Georgetown University (Protocol 2020-0033). Briefly, 5–6-week-old female NOD.Cg-*prkdc^scid^-IL2rg^tm1Wjl^/*SzJ (NSG) mice were purchased from Jackson Laboratory (JAX stock #005557) and housed at the Georgetown University animal facility. Mice were housed in a standard 12-h light-dark cycle and fed standard mouse feed and water ad libitum. Prior to xenografts, all mice were acclimated in standard housing conditions for a minimum of one week after delivery. For CDX first-generation studies, cells were harvested, washed with 1X PBS, and pelleted via centrifugation at 300x g for 4 minutes. Cells were resuspended in a 1:1 volume/volume solution of 1X DPBS (Life Technologies) and growth factor reduced Matrigel (Corning) up to a final total volume of 100 µl/injection. For second-generation CDX transplantation, cryopreserved 1mm^3^ tumor chunks in a 10% DMSO/90% FBS solution were thawed at room temperature and washed in 1X PBS twice prior to transplantation. Baseline weights were recorded for each animal and monitored throughout the study. Animals were anesthetized using 2-3% isoflurane, monitored for depth of anesthesia and respiration throughout the procedure. Surgical site was depilated and sterilized using ethanol and PVP Iodine prep swab sticks (PDI healthcare, cat. no. S41125) prior to xenografting. For first-generation CDXs cells were injected subcutaneously and for second-generation CDXs, a shallow incision was made, and pieces of first-generation CDX primary tumor no larger than 1mm^3^ were implanted in the unilateral dorsal flank of mice. Post operative wound closure, pain management, and monitoring was done as per the approved animal protocol. Animals were monitored weekly for the development of palpable tumors. Following palpable tumor formation, animals were monitored twice weekly and tumor size was measured via calipers. Tumor volume was calculated by the following formula: volume = 0.5 x L x W^2^ (where L is the length/longer dimension; W is the width/shorter dimension). Tumor-bearing mice were euthanized as per the animal protocol at specific time points depending on the experimental model as described in results. In general, mice were euthanized when the tumor reached 1500 mm^3^, 9 months post-injection, or when mice exhibited signs of poor health. Post euthanasia, primary tumors, metastases, and other organs were immediately harvested and sectioned for immunohistochemistry (IHC), cryopreservation in 10% DMSO:90%FBS (volume/volume), and flash freezing in liquid nitrogen for DNA/RNA extraction.

### RNA/DNA Extraction

RNA and DNA were isolated immediately from whole blood samples (“WBM”), healthy donor whole blood (“HD”), patient FFPE tissue biopsy cores (“FFPE”), CTC cultures (“TC”), and CDX organs. All RNA and DNA extraction kits were used following manufacturer’s protocols. All RNA samples were treated with RNAse-free DNAse provided by manufacturer’s kits or with RNase-Free DNAse Set (Qiagen, cat. No. 79254). All DNA samples were treated with RNAse provided by manufacturers to remove contaminant RNAs. FFPE Tissue cores of patient primary tumors or metastases were collected by the SRBSR, sectioned at the HTSR, and collected in microcentrifuge tubes. RNA and DNA were extracted from FFPE tissues via QIAamp DNA FFPE Tissue Kit (Qiagen, cat. no. 73504) or RNeasy FFPE Kit (Qiagen, cat. no. 56404). For TC samples, cultured cells were detached from culture plates by gently pipetting plates with 1X PBS or via mechanical removal with a cell scraper. Cells were pelleted via centrifugation (400x g for 4 minutes at 4°C) and washed one time with 1X PBS prior to RNA/DNA extraction. For TC samples, RNA was extracted using RNAqueous-Micro Total RNA Isolation Kits (Thermo Fisher, cat. no. AM1931) with added DNAse treatment. Resulting RNA was eluted in 13.5 µL of provided elution buffer. DNA from TC samples was extracted using either Qiagen DNeasy Blood & Tissue Kits (Qiagen, cat. no. 69504). DNA was eluted in 40 µL of ddH2O.

For CDX tissues, flash frozen tissue samples that had been prepared at the time of euthanasia were used. Tissues were first ground into a powder under liquid nitrogen freezing in mortar and pestles. RNA and DNA extraction proceeded immediately following grinding of CDX tissues. For DNA and RNA extraction, three kits were used: (1) Qiagen DNeasy Blood & Tissue Kits (Qiagen, cat. no. 69504) for DNA extraction and (2) RNeasy Mini Kit (Qiagen, cat. no. 74104) for RNA.

All DNA and RNA samples underwent an initial round of quality control via spectroscopy using a NanoDrop 2000 (Thermo Scientific). DNA samples were considered high quality with 260/280 ratios of ≥ 1.80. RNA samples were considered high quality with 260/280 ratios of ≥ 2.00. All DNA and RNA samples were immediately stored −80°C for long-term storage. Minimal freeze-thaw cycles were allowed to limit degradation.

### Immunohistochemistry and Immunofluorescence

Tissues collected from euthanized mice were fixed in formalin and paraffin embedded at the Histopathology & Tissue Shared Resource core at Georgetown University, American Histolabs Inc. at Gaithersburg, MD, or at the Yale Pathology Tissue Services at Yale University. Hematoxylin & Eosin of sectioned slides was performed at one of the three institutions, as well as at the Lombardi Comprehensive Cancer Center. Briefly, tissue sections were deparaffinized by melting at 60°C in an oven for 45 minutes followed by 2x xylene treatment for 20 minutes each. Slides were then rehydrated and antigen retrieval was done in manufacturer’s specified buffer (citrate buffer (pH 6.0), EDTA buffer (pH 8.0), or Tris/EDTA buffer (pH 9.0) made in house) at 97°C for 20 minutes in a PT module (Labvision, Kalamazoo, MI, USA). Endogenous peroxidase was blocked by using 0.3**%** hydrogen peroxide in methanol for 30 minutes in the dark followed by incubation of slides in a blocking buffer (0.3% bovine serum albumin in TBST (0.1 mol/L of TRIS-buffered saline (pH 7.0) containing 0.05% Tween-20)) for 30 minutes at room temperature. Slides were then incubated with primary antibody diluted in blocking buffer overnight at 4°C. Antibodies used in this study include anti-GAPDH (1:200 dilution, abcam, cat. no. ab128915), and anti-CD45 clone 2B11 + PD7/26 (Ready-to-use, Agilent Dako cat. no. GA75161-2). After washing away the primary antibodies, slides were incubated with secondary antibody (goat anti-mouse conjugated to horseradish peroxidase, supplied in Mouse and Rabbit Specific HRP/DAB (ABC) Detection IHC kit from Abcam cat. no. ab64264) to target either GAPDH or CD45 for one hour at room temperature. After washing, slides were treated with DAB substrate (abcam cat. no. ab64238) per manufacturer’s protocol. Immunohistochemistry for Vimentin was performed using IHCeasy Vimentin Ready-To-Use IHC kit (ProteinTech cat. no. KHC0039) following manufacturer’s protocol or an in-house standardized operating procedure. Control slides using tissue from a wildtype non-surgery NSG mice and human-specific tissue microarrays provided by the Yale Pathology Tissue Service were used to optimize DAB staining exposure. Exposure times were selected such that clear staining could be observed in positive human control slides, but not in uninjected mouse slides. No exposure lasted longer than 10 minutes. Following DAB staining, slides were washed two times with ddH2O and counterstained using hematoxylin solution (Abcam cat. no. ab220365) or counterstain supplied in kit for 1-2 minutes. Slides were subsequently washed twice in water to remove excess counterstain and covered using glass coverslips secured with Fluorsave Aqueous Mounting Medium (Sigma-Aldrich cat. no. 345789). If coverslips needed to be removed, slides were soaked overnight in ddH2O at 4°C.

For EpCAM immunofluorescence, slides were incubated with primary antibody, anti-EpCAM (1:200 dilution, abcam cat. no. ab223582), diluted in blocking buffer overnight at 4°C. After washing away the primary antibodies, slides were incubated with secondary antibody (Donkey-anti-mouse conjugated with Alexa Fluor 568, Invitrogen, cat. no. A21202) for one hour at room temperature. After incubation, slides were washed 2X with TBST and 1X with TBS, and incubated for 10 additional minutes using Hoescht 33342 Solution (Thermo Fisher, cat. no. 62249) per manufacturer’s protocol. Slides were then washed 2X with TBST and 1X with TBS and mounted using either Fluorsave Aqueous Mounting Medium or ProLong Gold Antifade Reagent with DAPI (Thermo Fisher, cat. no. P36935).

### Quantitative Real-Time PCR

Following RNA extraction, RNA was converted to cDNA using LunaScript RT SuperMix Kit (NEB, cat. no. E3010) according to manufacturer specifications. For all reactions, a minimum of 25 µg and maximum of 100 µg of RNA was used as input, with a final reaction volume of 20 µL. A Bio-Rad T100 thermal cycler was used to incubate reactions using the following program: 25°C for 2 minutes; 55°C for 10 minutes; 95°C for 1 minute; 4°C hold. All cDNA reactions were subsequently diluted with ddH2O to a final concentration of 25 µg initial input RNA/20 µL. All cDNA were stored at −20°C for long-term storage, with minimal freeze-thaw cycles allowed to limit degradation.

For quantitative RT-PCR, Luna Universal qPCR MasterMix (NEB, cat. no. M3003) was used according to manufacturer’s specifications. Briefly, 20 µL reactions were mixed using 10 µL of Luna Universal qPCR Master Mix, 0.5 µL per Forward and Reverse Primer (diluted to 10 µM), 1 µL of cDNA template, and nuclease-free water to final reaction volume. PCR primers (provided in 5’ to 3’ format). All qRT-PCR reactions were run in a Bio-Rad CFX 96 well thermal cycler with Thermo Scientific PCR 96-well plates (Thermo Fisher, cat. no. AB-0800). All qRT-PCR reactions were analyzed using the 2^-ΔΔCt^ method. All qRT-PCR samples were run through gel electrophoresis using in-house made 1.5% agarose gels run at 95V for 35 minutes prior to imaging. All electrophoresis images were taken using a GE AI600 RGB Gel imaging system.

### Next-generation RNA and Whole Exome Sequencing Library Preparation and Sequencing

All next generation sequencing library preparation and sequencing was performed with the support of established third-party commercial companies. Library preparation and RNA-sequencing was conducted by Psomagen (Rockville, MD). Libraries were created using TruSeq stranded mRNA sample preparation and paired-end reads with a targeted read-length of 151 bp were performed on a NovaSeq 6000 S4 sequencing platform. Low-input RNA-sequencing was performed, requiring a minimum of 50 ng of RNA for processing. For each sample, 60M total minimum reads was targeted; for CDX samples with potential for mouse cell contamination, 100M total reads was targeted. For whole exome sequencing, libraries were created using SureSelect V5-post library preparation kits and paired-end reads with a targeted read-length of 151 bp were sequenced on a NovaSeq 6000 S4 sequencing platform. A minimum of 500 ng of high-quality DNA was required for library preparation and sequencing. For each sample, 100X raw depth coverage was targeted.

### Bulk RNA-sequencing Bioinformatics Analysis

Paired-end read files were trimmed using Trimmomatic^75^ and aligned to the human transcriptome (GRCh38.p13) reference from Gencode v33 using salmon v0.14. The data quality was verified using FastQC. The salmon output was processed in R using the tximport and tidyverse packages^76^. The differential expression analysis was performed in R using DESeq2^77^. Normalized feature counts were used with the gene set enrichment analysis, GSEA software and Molecular Signature Database (MSigDB), available at http://www.broad.mit.edu/gsea/44. GSEA analysis was performed in R using the open-source package fgsea (version 1.20.1)^78^. Genesets for fgsea input were downloaded from the MSigDB database^44^. KEGG analysis was performed in R using the open-source package clusterProfiler (Release 4.2.1)^79^. Other R source packages used for figure development include but are not limited to ggsci and gplots as well as Matlab R2021a/b. CIBERSORT was accessed via https://www.cibersort.stanford.edu/ and non-normalized feature counts were input following developer instructions^37^.

### Whole Exome-Sequencing Bioinformatics Analysis

To identify mutations and genetic alterations associated with metastatic phenotypes, whole exome sequencing of cultured CTCs and corresponding patient FFPE tissue and healthy donors was performed. Briefly, deep sequencing and exome capture was performed using SureSelect V5-post and Illumina platform with 200X coverage. Using BWA (v0.7.17), we indexed human (hg38) and mouse (mm10) chromosomal sequences (fasta format). Then BWA mem2 module was then used to align raw FASTQ files to the human reference genome (GRCh38.p13). The contaminating mouse sequences were removed using the XenofilteR package in R. The package estimates edit distance to classify sequences as mouse or human and then removes the mouse sequences.

The resulting bam (mapped to hg38 or xeno-filtered) files were GATK and SnpEff. As a result, were generated for predicted SNP and InDel events and this information was used for final interpretations in this manuscript. For reproducibility purposes, we are maintaining our WES scripts in this repository https://github.com/goodarzilab/WES. Mutant-allele tumor heterogeneity (MATH) scores were generated for CTC models and compared to patient’s FFPE. Somatic mutations, consisting of point mutations, insertions, and deletions across the exome were identified using the VariantDx custom software^80^. Somatic mutations were annotated against the set of mutations in COSMIC (v.84) database. Revel scores for missense mutations were calculated to determine mutation potential as cancer drivers by CHASMplus^81^. CTC or FFPE exclusive mutations were defined based on a MAF ≥ 0.5 and ≤ 0.25 in the opposing corresponding sample type from the same patient. Shared mutations were defined based on a MAF ≥ 0.5 in all samples. Only SNVs with a mutation count ≥ 4 were considered real and included within the differential enrichment analysis.

## Funding and Acknowledgements

This study was funded in part by a research grant from the Ruesch Center for the Cure of Gastrointestinal Cancers, Georgetown University and support from the Center for Cell Reprogramming at Georgetown University to SA. Illustrations were created with the help of Biorender.com. This research was supported by the SRBSR, HTSR and DCM funded by the NIH Cancer Center Support Grant to Georgetown Lombardi Comprehensive Cancer Center (P30-CA051008). JX was partly supported by a Medical Graduate Student Organization (MCGSO) mini-grant.

## Ethics approval and consent to participate

Animal experiments with mice were approved by the Georgetown University Institutional Animal Care and Use Committee, Protocol #2020-0033.

### Consent for publication

All authors have reviewed the data and agreed on submission of the manuscript.

## Author Contributions

**Conception and Design**: S. Agarwal, R, Schlegel, Paula R. Pohlmann

**Development and Optimization of Methodology:** J. Xiao, S. Agarwal

**Acquisition of Data**: J. Xiao, U. Sharma, S. Suguru

**Computational Analysis:** H. Goodarzi, S. Miglani, A. Arab, S. Bhalla, R. Suter

**Recruitment of Patients, Acquisition of Blood Samples and Clinical Information:** R. Mukherji, P. R. Pohlmann, J. Marshall, B. Weinberg, A. R. He, M. Noel

**Data Interpretation**: J. Xiao, S. Agarwal, R. Schlegel, M. Lippman

**Writing of the Manuscript**: Initial draft is written by J. Xiao, S. Agarwal

**Editing of Manuscript:** All authors participated in editing the manuscript

**Study Supervision:** S. Agarwal

**Data and materials availability**: RNA and DNA sequencing data in the current study have been deposited in the Gene Expression Omnibus (GEO).

## SUPPLEMENTAL FIGURES/TABLES

**Figure S1.**
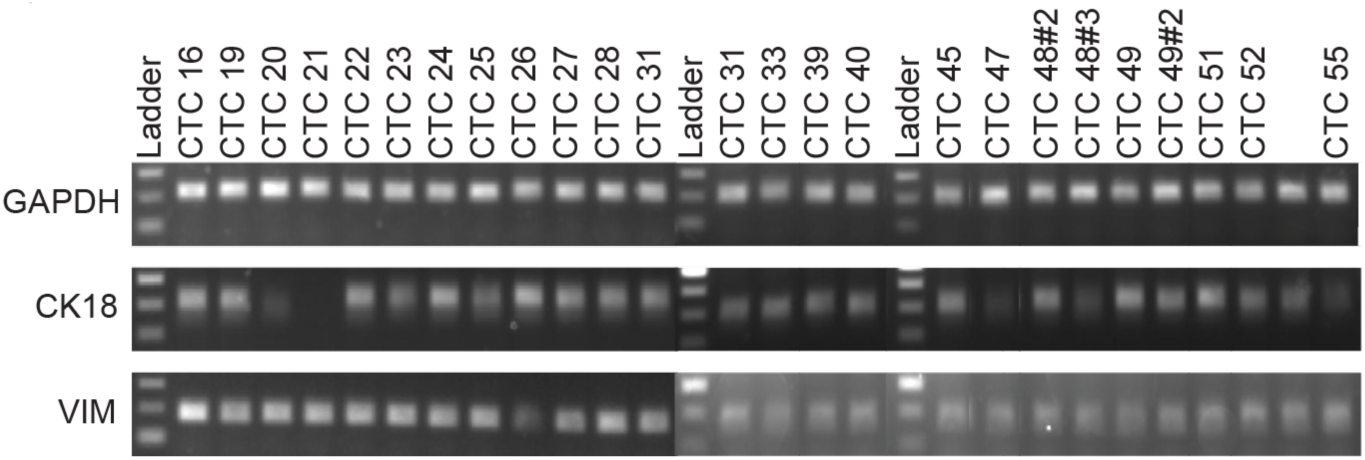
qRT-PCR was performed to confirm expression of human-specific cytokeratin 18 and vimentin in various cultured CTCs.

**Figure S2.**
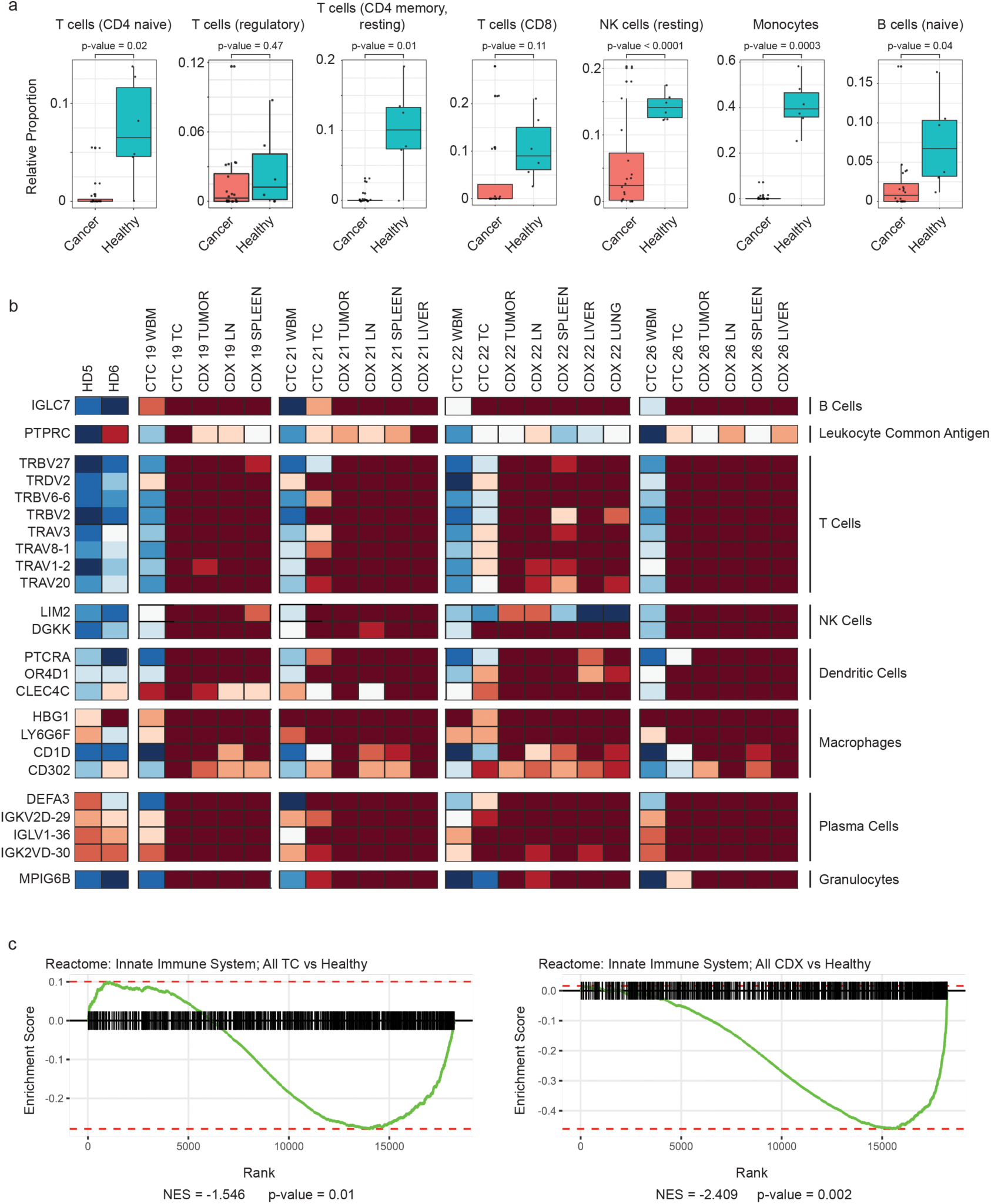
(a) Box plots of the relative proportion of immune cells signatures from CIBERSORT deconvolution indicates a depletion of T and B cells, as well as natural killer (NK) cells and monocytes. (b) Panels of genes that are highly enriched in single-cell immune signatures from Protein Atlas were extracted from the RNA-sequencing, confirming a depletion of immune cells within cancer samples except for corresponding patient whole blood. (c) Reactome pathway analysis was performed in comparisons between CTC culture (TC) or CDX samples against healthy. Both analyses indicate a depletion of innate immune cell system genes in either sample type.

**Figure S3.**
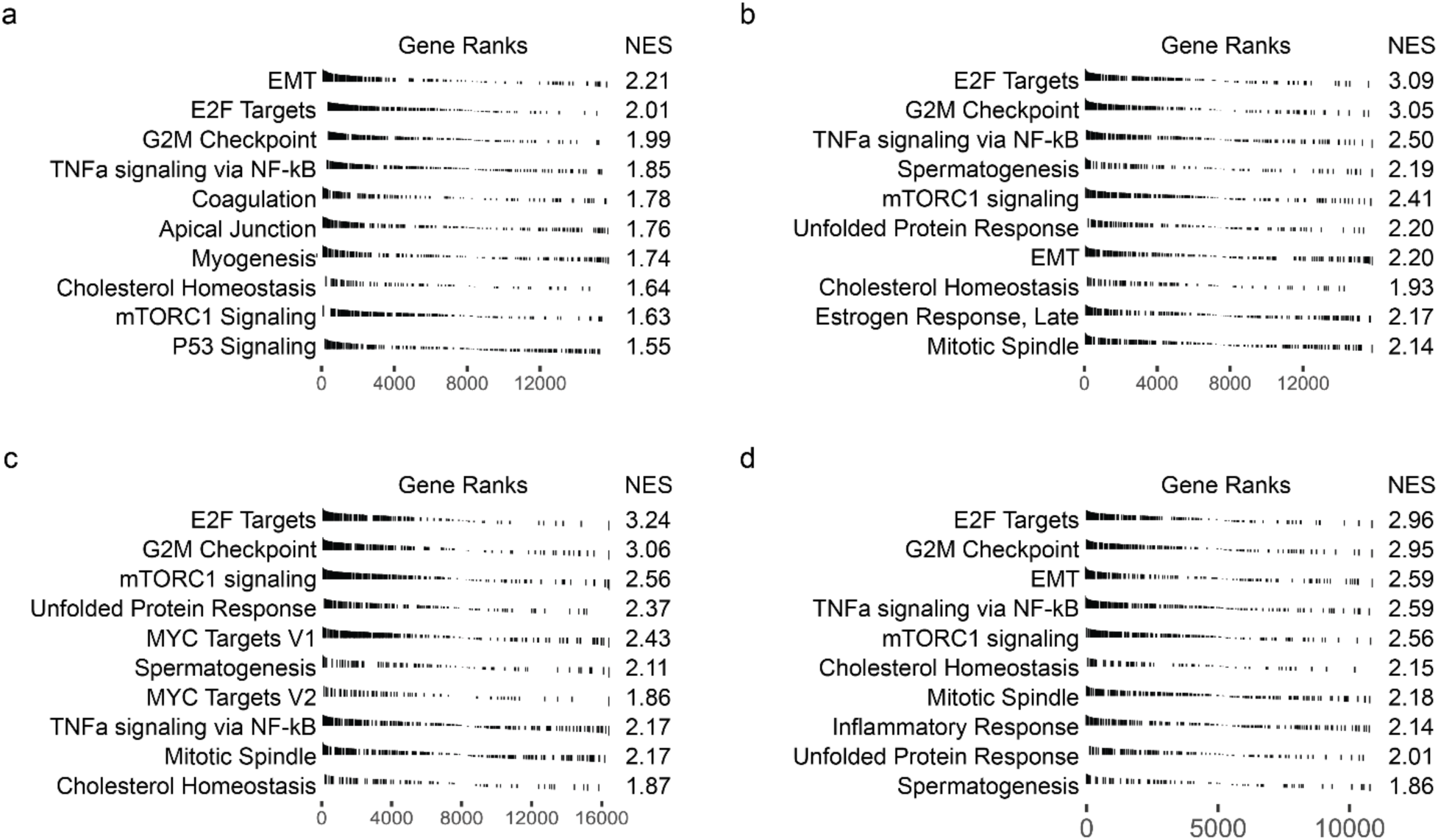
GSEA analysis was performed to evaluate enrichment in all CTC-derived samples relative to whole blood samples in (a) patient #19, (b) patient #21, (c) patient #22, and (d) patient #26.

**Figure S4.**
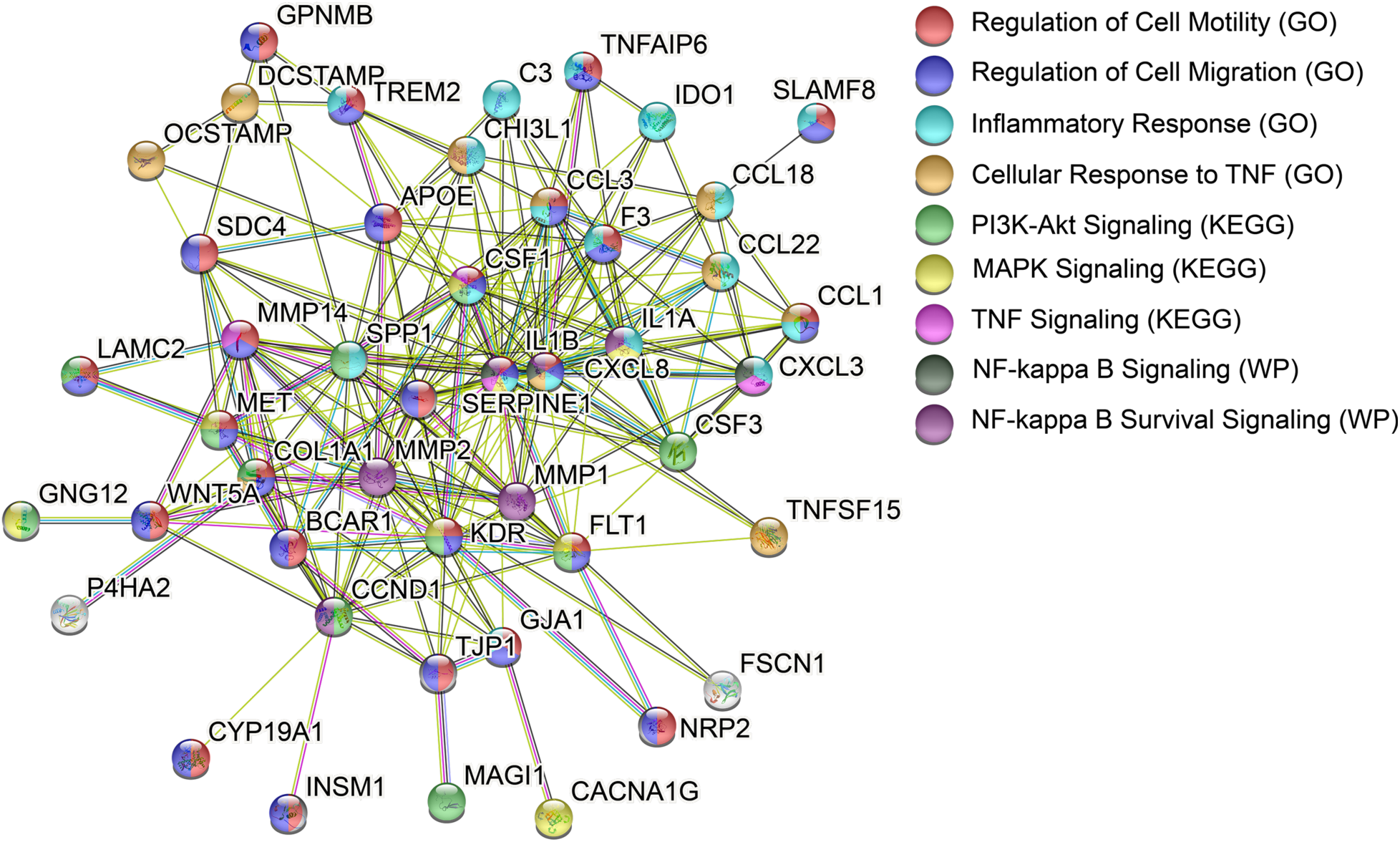
Functional network analysis reveals potential interactome circuitry unique to CTC cultures. A functional network was generated through input of the 199 gene prospective CTC culture signature in Table S3 into String-DB, using the whole human genome as background. Nodes represent all proteins produced by a single, protein-coding gene locus, and are colored by associated gene ontology (GO), KEGG pathway, or WikiPathway (WP) terms. White nodes represent proteins in a second shell of interactors. Edges or connecting lines represent protein-protein interactions. Known interactions are depicted as teal (from curated databases) or magenta (experimentally determined) edges. Predicted interactions are depicted as green (gene neighborhood), red (gene fusions), blue (gene co-occurrence), yellow (from text mining), black (co-expression), or violet (protein homology). Disconnected nodes were not plotted. All enriched terms showed a FDR < 0.05.

**Table S1.** Differential gene analysis summarizing various cancer-agnostic and patient-specific comparisons.

| Sample | Control | Upregulated Genes | Downregulated Genes |
| --- | --- | --- | --- |
| All CTC-derived samples | Whole blood samples | 848 | 21 |
| Cultured CTCs | Whole blood samples | 1828 | 1165 |
| CDX Tissues | Whole blood samples | 3129 | 3419 |
| Cultured CTCs | CDX Tissues | 1836 | 2731 |
| **CTC 19 Specific Analysis** |  |  |  |
| All CTC-derived samples | Whole blood samples | 213 | 12 |
| Cultured CTCs | Whole blood samples | 290 | 9463 |
| CDX Tissues | Whole blood samples | 2586 | 3122 |
| Cultured CTCs | CDX Tissues | 9279 | 191 |
| **CTC 21 Specific Analysis** |  |  |  |
| All CTC-derived samples | Whole blood samples | 688 | 187 |
| Cultured CTCs | Whole blood samples | 474 | 656 |
| CDX Tissues | Whole blood samples | 1604 | 2685 |
| Cultured CTCs | CDX Tissues | 244 | 626 |
| **CTC 22 Specific Analysis** |  |  |  |
| All CTC-derived samples | Whole blood samples | 796 | 256 |
| Cultured CTCs | Whole blood samples | 1533 | 791 |
| CDX Tissues | Whole blood samples | 1970 | 3531 |
| Cultured CTCs | CDX Tissues | 733 | 2602 |
| **CTC 26 Specific Analysis** |  |  |  |
| All CTC-derived samples | Whole blood samples | 359 | 65 |
| Cultured CTCs | Whole blood samples | 1138 | 1297 |
| CDX Tissues | Whole blood samples | 1570 | 3011 |
| Cultured CTCs | CDX Tissues | 1323 | 2322 |

**Table S2.** Top 10 most up- and down-regulated genes, patient-specific.

| Upregulated (Gene, log2-fold change) |  |  |  |
| --- | --- | --- | --- |
| CTC 19 | CTC 21 | CTC 22 | CTC 26 |
| GDA (25.8) | FOXA2 (24.5) | GJB1 (22.9) | NR2F2 (25.7) |
| IVL (25.7) | DSC3 (24.4) | NES (22.6) | CCN1 (25.0) |
| CCL17 (25.6) | MT1G (24.3) | FGA (21.8) | CCDC74A (13.4) |
| SYNPO2L (24.7) | IVL (23.6) | FLRT2 (21.2) | CERNA2 (12.2) |
| FRMPD2 (24.4) | HOXD9 (23.5) | GAL (20.7) | CXCL13 (11.7) |
| KCNA1 (24.3) | MT1M (23.4) | COL3A1 (20.0) | UCHL1 (11.5) |
| SIX3 (23.7) | GJB1 (23.1) | CHRD (19.9) | HMX2 (11.1) |
| GRIN2A (23.1) | PDPN (23.06) | S100A16 (19.9) | INHBA (11.0) |
| CCDC74A (13.0) | RASD2 (22.6) | OVOL1 (19.7) | VSTM2L (10.9) |
| BCAR1 (12.6) | FLRT2 (22.6) | CALB2 (19.7) | EFNB2 (10.7) |
| Downregulated (Gene, log2-fold change) |  |  |  |
| XIST (-28.2) | XIST (-28.5) | RPS4Y1 (-30.0) | RPS4Y1 (-30) |
| CLEC4OP (-7.9) | SIGLEC14 (-9.4) | EIF1AY (-27.0) | EIF1AY (-26.4) |
| IGHV7-4-1 (-7.9) | ZFP57 (-7.3) | TXLNGY (-26.6) | TXLNGY (-25.8) |
| HDC (-7.8) | NRGN (-6.4) | KDM5D (-26.2) | UTY (-25.7) |
| SPATC1L (-7.7) | GZMH (-6.2) | DDX3Y (-25.6) | HSPA7 (-9.2) |
| IGLC5 (-6.8) | GZMM (-6.1) | GSTM1 (-10.4) | ZFP57 (-8.2) |
| LYPD2 (-6.6) | PTCRA (-6.1) | UTY (-10.1) | GSTM1 (-8.0) |
| MS4A2 (-6.4) | TRDC (-6.1) | LYPD2 (-8.0) | LCN8 (-7.7) |
| VMO1 (-5.6) | TMEM40 (-5.9) | LCN8 -7.6) | KRT72 (-7.3) |
| FCER1A (-5.6) | ITGA2B (-5.9) | PRKY (-7.2) | RNASE3 (-7.1) |

**Table S3.** A prospective CTC culture signature based on enriched expression in CTC cultures relative to patient-matched controls.

|  |  |  |  |  |
| --- | --- | --- | --- | --- |
| A4GALT | CP | HSD17B14 | NRP2 | SPR |
| ADAMDEC1 | CRABP2 | HSPB7 | OCSTAMP | SPRY4 |
| ADRA2B | CSF1 | IDO1 | OVOL1 | SRRM3 |
| AGRN | CSF3 | IGSF3 | P4HA2 | SSTR2 |
| AHRR | CSPG4 | IL1A | PAQR5 | ST18 |
| AK4 | CTSK | IL1B | PCARE | STC2 |
| ANKRD1 | CXCL3 | IL4I1 | PCGF2 | STEAP1 |
| AOC1 | CXCL8 | INHBA-AS1 | PDCD1LG2 | STRA6 |
| AOX1 | CYP19A1 | INSM1 | PDLIM4 | SVEP1 |
| APOC1 | CYP27B1 | KDR | PHF24 | SYT12 |
| APOE | DCSTAMP | KLK4 | PIR | TCEAL9 |
| ARMC9 | DEPP1 | KRT78 | PLOD2 | TFPI2 |
| ASPHD1 | DGKI | KRT79 | PLTP | TGM2 |
| ATF3 | DIRAS2 | LAMC2 | PNCK | TIE1 |
| BCAN-AS1 | DPYSL3 | LGMN | PNPLA3 | TJP1 |
| BCAR1 | DUSP4 | LINC00520 | PPIC | TM4SF1 |
| BEND3P2 | EBI3 | LINC01010 | RAB42 | TMEM132A |
| BHLHE41 | EMP1 | LINC01283 | RAB7B | TMEM200B |
| C15orf48 | EMX1 | LINC01426 | RAI14 | TMEM45A |
| C1QTNF1 | ENTHD1 | LINC01827 | RASAL2 | TMEM51-AS1 |
| C2CD4B | EPB41L1 | LINC02099 | RBP1 | TNFAIP6 |
| C3 | EPOP | LINC02137 | RGS16 | TNFAIP8L3 |
| C6orf223 | F3 | LINC02244 | RGS20 | TNFSF15 |
| CA12 | FAM124A | LIPG | RHBDF1 | TREM2 |
| CACNA1G | FJX1 | LRRC32 | RN7SL368P | TRIM6 |
| CAVIN1 | FLRT2 | MAGI1 | RNASE1 | TSKU |
| CCDC74A | FLT1 | MET | RPLP0P2 | TTC23 |
| CCL1 | FSCN1 | MMP1 | RRAD | UST-AS2 |
| CCL18 | GAL | MMP14 | RTN4RL2 | VWA1 |
| CCL22 | GDA | MMP2 | SCG5 | WNT5A |
| CCL3 | GDF15 | MRC1 | SDC2 | ZMIZ1-AS1 |
| CCM2L | GEM | MS4A6E | SDC4 |  |
| CCND1 | GJA1 | MT1G | SERPINE1 |  |
| CD109 | GJA5 | MT1H | SEZ6L2 |  |
| CD276 | GJB2 | MT1L | SHC4 |  |
| CD83 | GNG12 | MT1M | SLAMF8 |  |
| CHI3L1 | GPC4 | MUCL1 | SLAMF9 |  |
| CHST3 | GPNMB | MYPN | SLC1A2 |  |
| CLCNKA | GPX3 | NCS1 | SLC41A2 |  |
| CLLU1-AS1 | GSDME | NECTIN4 | SLC7A11 |  |
| COL1A1 | HRH1 | NES | SLCO2B1 |  |
| COL22A1 | HSD11B1 | NPR1 | SPP1 |  |

**Table S4.** A prospective CDX signature based on enriched expression in CDX primary tumor samples.

|  |  |  |  |  |
| --- | --- | --- | --- | --- |
| ABCA12 | CXCL9 | KCNN1 | PIEZO2 | TFPI2 |
| ADAMTS12 | DMD | KIAA1549L | POU2F3 | TSPAN11 |
| BARX2 | EMX1 | LHX2 | PRELID1P5 | TSPAN12 |
| BCAR1 | ENOX1 | LINC01055 | RHOV |  |
| CCL1 | GJC1 | LINC01258 | RRAD |  |
| CCL22 | HNRNPA1P30 | LINC02227 | SLC12A8 |  |
| CCN1 | IGSF3 | LSAMP | SYNPO2L |  |
| COL1A1 | IL13 | OSR2 | TBXT |  |

**Table S5.** TCGA pan-cancer atlas was queried to identify genes whose expression correlated with poorer overall survival.

| Gene | Hazard Ratio | P-value |
| --- | --- | --- |
| BCAR1 | 1.2 (1.0 - 1.5) | 0.027 |
| COL1A1 | 1.6 (1.3 - 2.0) | < 0.001 |
| IGSF3 | 1.3 (1.1 - 1.6) | 0.005 |
| RRAD | 1.4 (1.1 - 1.7) | 0.002 |
| TFPI2 | 1.5 (1.2 - 1.8) | < 0.001 |
| CCL22 | 1.1 (0.91 - 1.4) | 0.278 |
| EMX1 | 0.98 (0.8 - 1.2) | 0.807 |

**Table S6.**
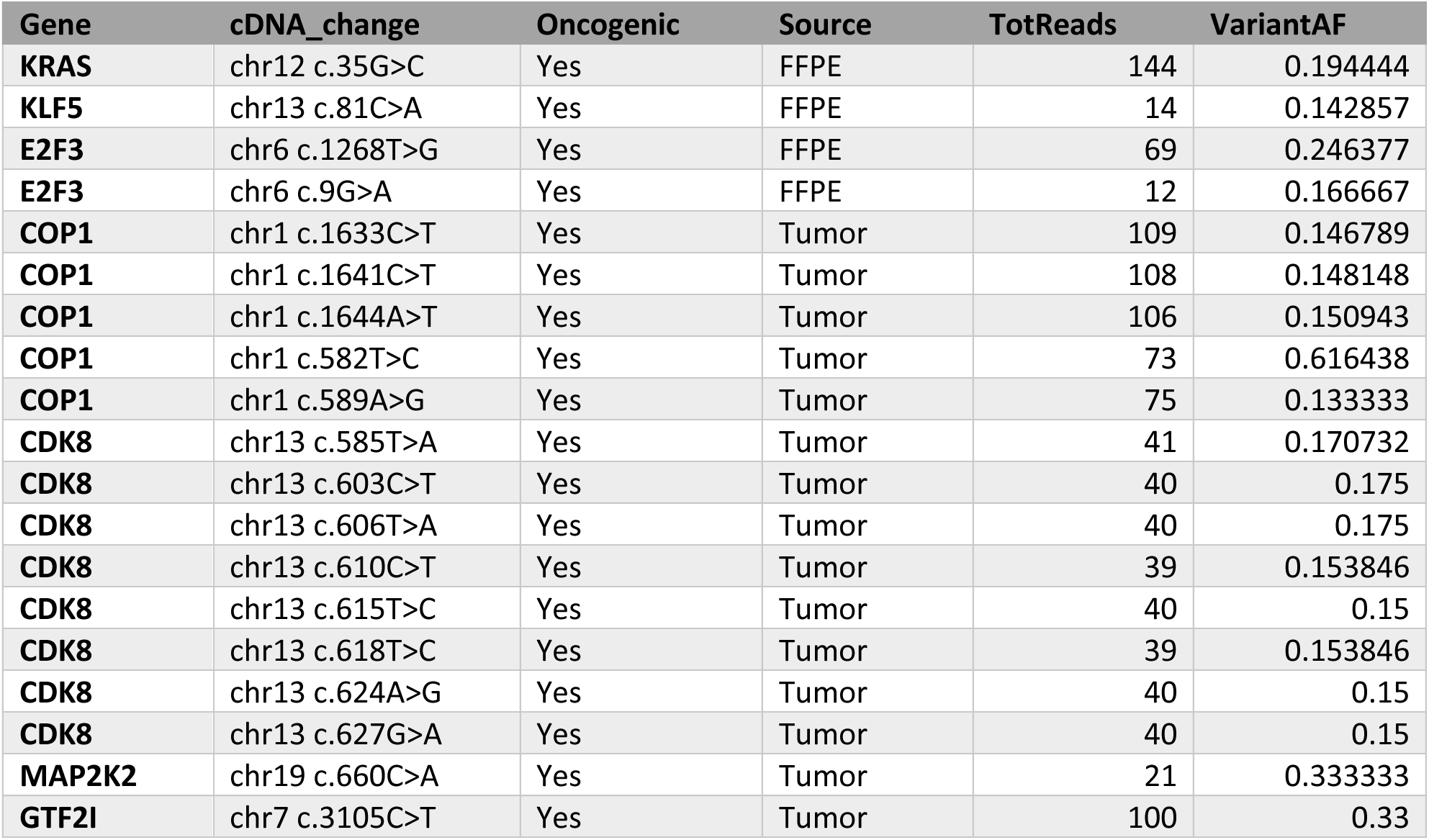
Oncogenic mutations that were either FFPE, CTC culture, or CDX tumor-exclusive in Patient #19.

| Gene | cDNA_change | Oncogenic | Source | TotReads | VariantAF |
| --- | --- | --- | --- | --- | --- |
| KRAS | chr12 c.35G>C | Yes | FFPE | 144 | 0.194444 |
| KLF5 | chr13 c.81C>A | Yes | FFPE | 14 | 0.142857 |
| E2F3 | chr6 c.1268T>G | Yes | FFPE | 69 | 0.246377 |
| E2F3 | chr6 c.9G>A | Yes | FFPE | 12 | 0.166667 |
| COP1 | chr1 c.1633C>T | Yes | Tumor | 109 | 0.146789 |
| COP1 | chr1 c.1641C>T | Yes | Tumor | 108 | 0.148148 |
| COP1 | chr1 c.1644A>T | Yes | Tumor | 106 | 0.150943 |
| COP1 | chr1 c.582T>C | Yes | Tumor | 73 | 0.616438 |
| COP1 | chr1 c.589A>G | Yes | Tumor | 75 | 0.133333 |
| CDK8 | chr13 c.585T>A | Yes | Tumor | 41 | 0.170732 |
| CDK8 | chr13 c.603C>T | Yes | Tumor | 40 | 0.175 |
| CDK8 | chr13 c.606T>A | Yes | Tumor | 40 | 0.175 |
| CDK8 | chr13 c.610C>T | Yes | Tumor | 39 | 0.153846 |
| CDK8 | chr13 c.615T>C | Yes | Tumor | 40 | 0.15 |
| CDK8 | chr13 c.618T>C | Yes | Tumor | 39 | 0.153846 |
| CDK8 | chr13 c.624A>G | Yes | Tumor | 40 | 0.15 |
| CDK8 | chr13 c.627G>A | Yes | Tumor | 40 | 0.15 |
| MAP2K2 | chr19 c.660C>A | Yes | Tumor | 21 | 0.333333 |
| GTF2I | chr7 c.3105C>T | Yes | Tumor | 100 | 0.33 |

**Table S7.**
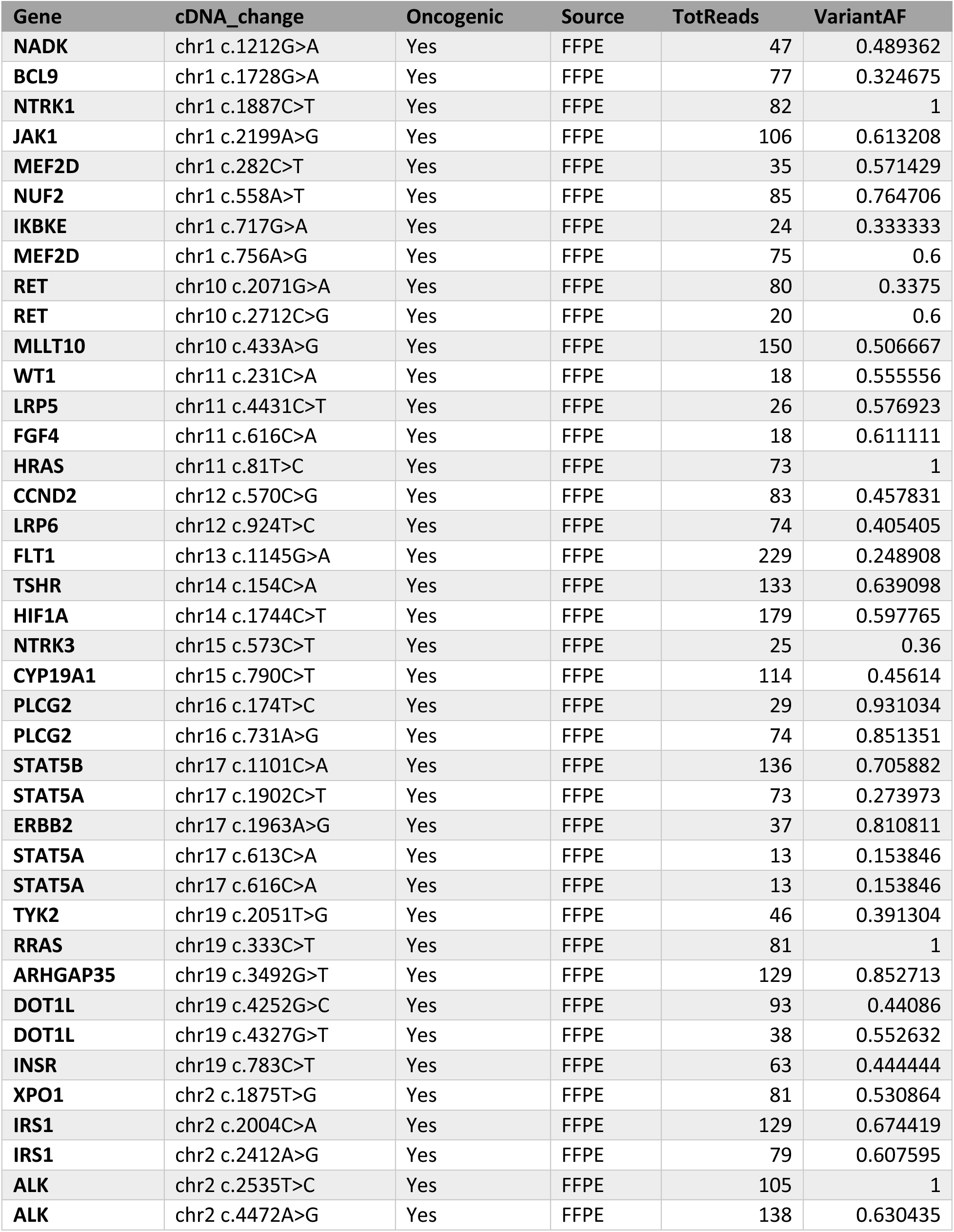
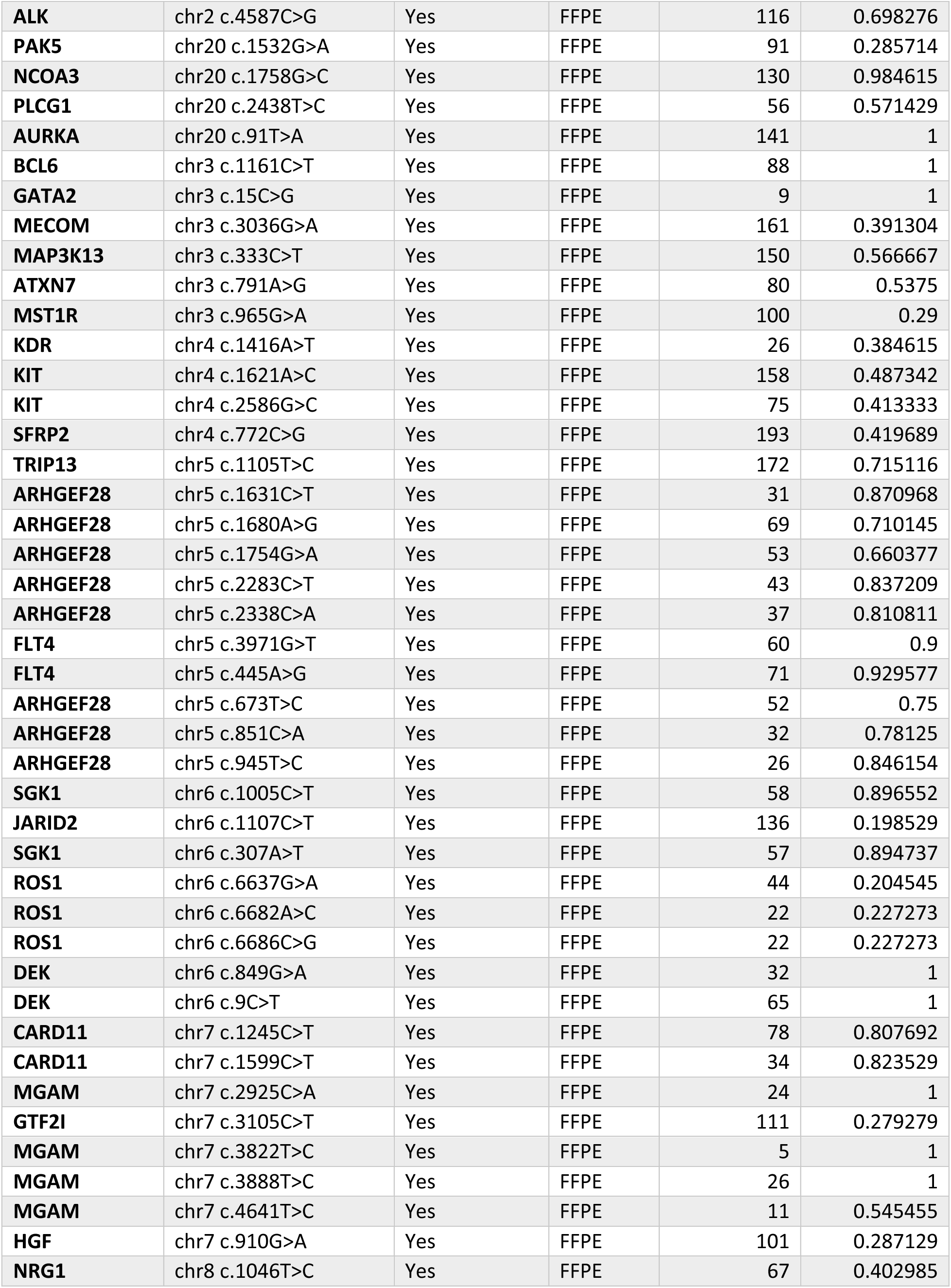
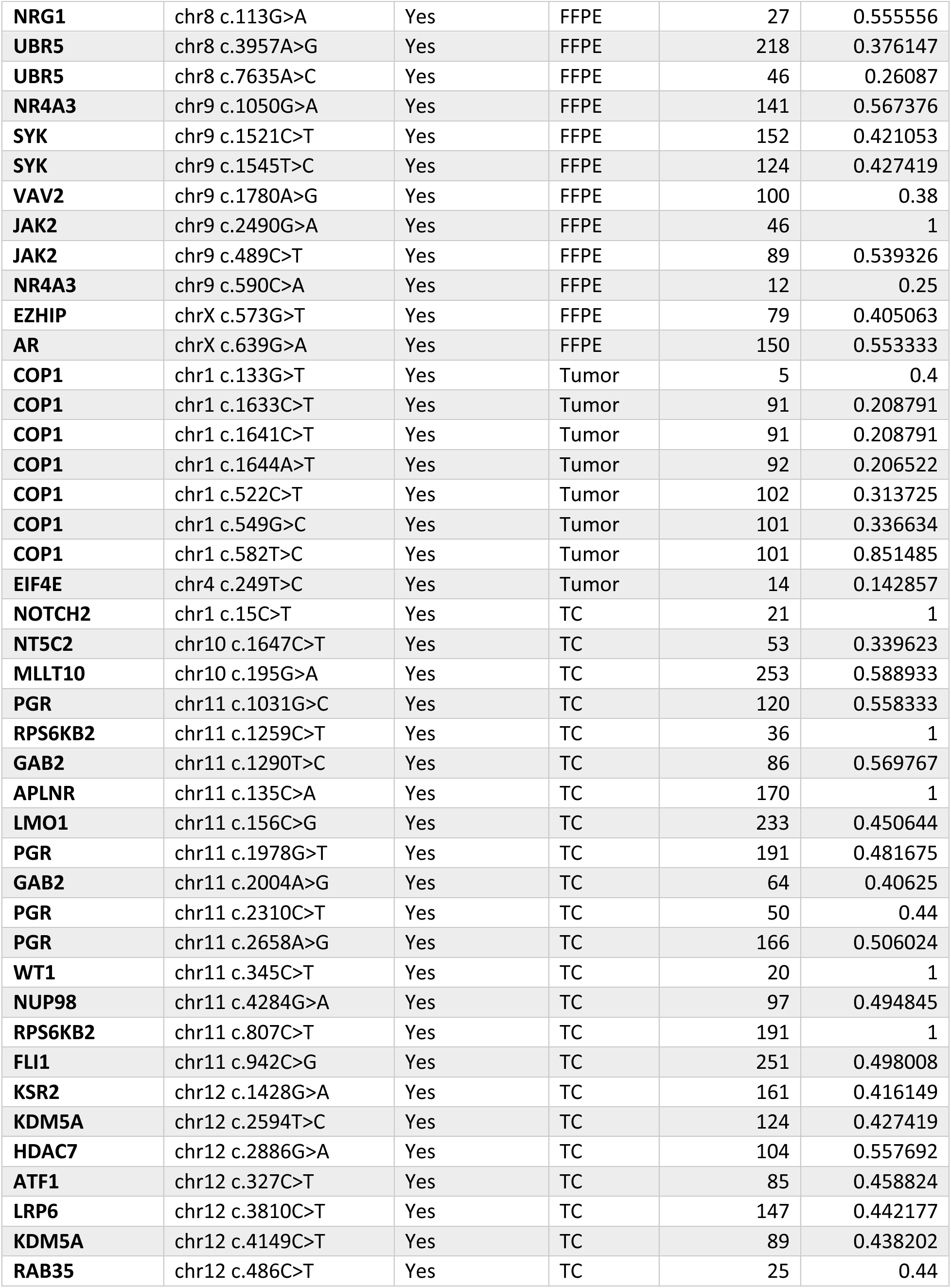
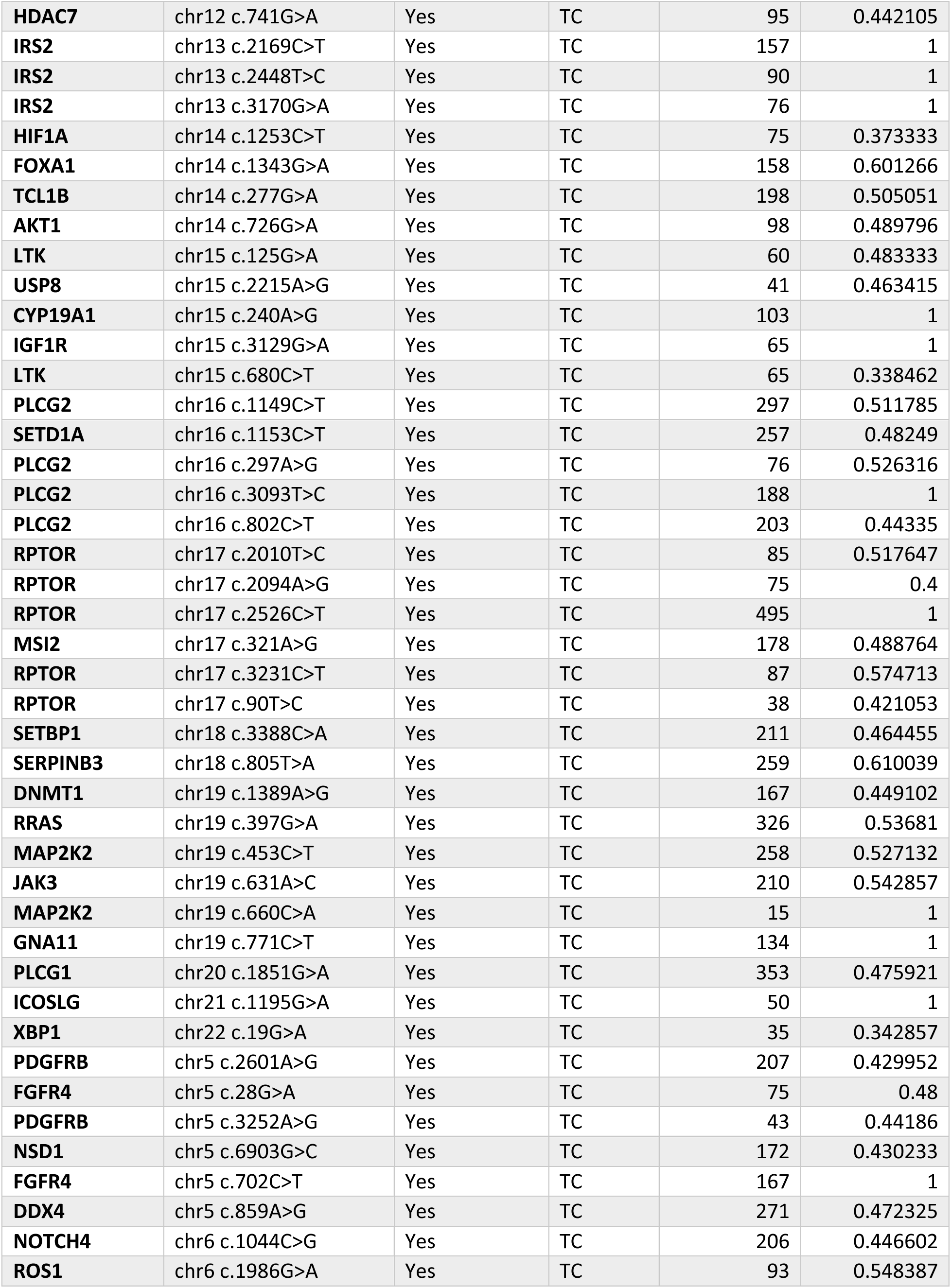
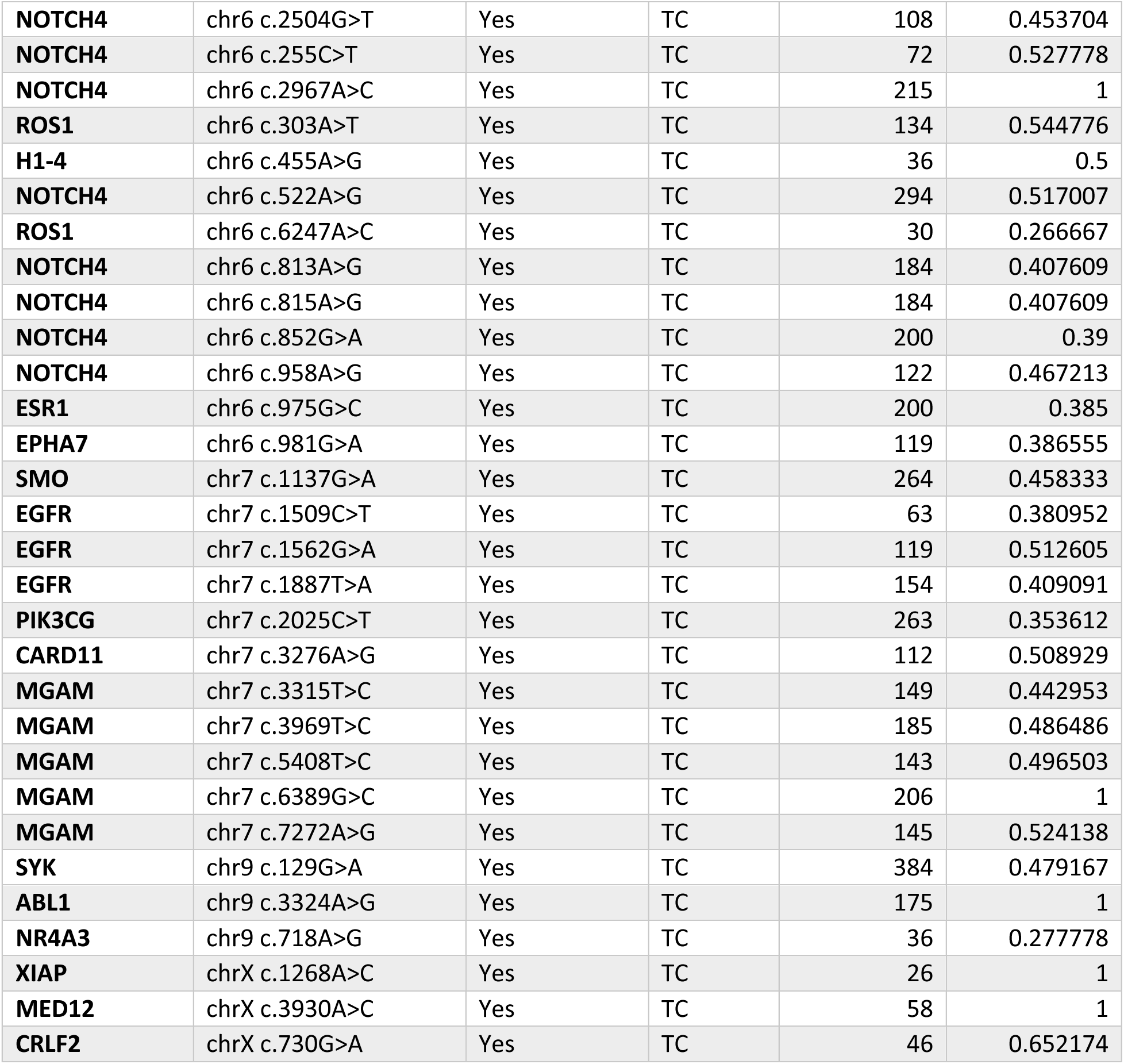
Oncogenic mutations that were either FFPE, CTC culture, or CDX tumor-exclusive in Patient #21.

**Table S8.** Oncogenic mutations that were either FFPE, CTC culture, or CDX tumor-exclusive in Patient #22.

| Gene | cDNA_change | Oncogenic | Source | TotReads | VariantAF |
| --- | --- | --- | --- | --- | --- |
| **FLT1** | chr13 c.1145G>A | Yes | FFPE | 229 | 0.248908 |
| **STAT5A** | chr17 c.613C>A | Yes | FFPE | 13 | 0.153846 |
| **STAT5A** | chr17 c.616C>A | Yes | FFPE | 13 | 0.153846 |
| **JARID2** | chr6 c.1107C>T | Yes | FFPE | 136 | 0.198529 |
| **MGAM** | chr7 c.3822T>C | Yes | FFPE | 5 | 1 |
| **NR4A3** | chr9 c.590C>A | Yes | FFPE | 12 | 0.25 |
| **COP1** | chr1 c.582T>C | Yes | Tumor | 21 | 0.571429 |
| **GNAS** | chr20 c.636G>A | Yes | Tumor | 8 | 0.5 |
| **NUF2** | chr1 c.686C>T | Yes | TC | 22 | 0.409091 |

**Table S9.** Oncogenic mutations that were either FFPE, CTC culture, or CDX tumor-exclusive in Patient #26.

| Gene | cDNA_change | Oncogenic | Source | TotReads | VariantAF |
| --- | --- | --- | --- | --- | --- |
| PIK3CD | chr1 c.2292C>T | Yes | FFPE | 6 | 0.333333 |
| GNB1 | chr1 c.908A>G | Yes | FFPE | 2 | 1 |
| LRP6 | chr12 c.499G>A | Yes | FFPE | 2 | 1 |
| SOX9 | chr17 c.1328C>T | Yes | FFPE | 2 | 1 |
| HOXB13 | chr17 c.249T>C | Yes | FFPE | 2 | 1 |
| YES1 | chr18 c.778C>A | Yes | FFPE | 3 | 0.666667 |
| NOTCH3 | chr19 c.1736G>C | Yes | FFPE | 3 | 0.666667 |
| NOTCH3 | chr19 c.1744C>A | Yes | FFPE | 3 | 0.666667 |
| IRS1 | chr2 c.3073A>G | Yes | FFPE | 3 | 0.666667 |
| NSD2 | chr4 c.2909G>A | Yes | FFPE | 255 | 0.164706 |
| NOTCH4 | chr6 c.1794T>C | Yes | FFPE | 2 | 1 |
| COP1 | chr1 c.1633C>T | Yes | Tumor | 85 | 0.188235 |
| COP1 | chr1 c.1641C>T | Yes | Tumor | 85 | 0.188235 |
| COP1 | chr1 c.1644A>T | Yes | Tumor | 85 | 0.188235 |
| COP1 | chr1 c.1650A>G | Yes | Tumor | 82 | 0.170732 |
| COP1 | chr1 c.428T>C | Yes | Tumor | 15 | 0.2 |
| COP1 | chr1 c.432A>G | Yes | Tumor | 15 | 0.333333 |
| COP1 | chr1 c.435A>G | Yes | Tumor | 13 | 0.230769 |
| COP1 | chr1 c.438A>C | Yes | Tumor | 15 | 0.133333 |
| COP1 | chr1 c.522C>T | Yes | Tumor | 80 | 0.3125 |
| COP1 | chr1 c.549G>C | Yes | Tumor | 81 | 0.308642 |
| COP1 | chr1 c.582T>C | Yes | Tumor | 84 | 0.809524 |
| COP1 | chr1 c.591G>A | Yes | Tumor | 88 | 0.25 |
| COP1 | chr1 c.615C>A | Yes | Tumor | 50 | 0.36 |
| FGFR2 | chr10 c.1709G>T | Yes | Tumor | 13 | 0.153846 |
| LMO2 | chr11 c.100G>T | Yes | Tumor | 8 | 0.25 |
| IRS2 | chr13 c.2851G>A | Yes | Tumor | 11 | 0.181818 |
| CDK8 | chr13 c.324C>T | Yes | Tumor | 53 | 0.188679 |
| CDK8 | chr13 c.342T>A | Yes | Tumor | 82 | 0.134146 |
| CDK8 | chr13 c.348A>C | Yes | Tumor | 87 | 0.126437 |
| CDK8 | chr13 c.585T>A | Yes | Tumor | 29 | 0.310345 |
| CDK8 | chr13 c.603C>T | Yes | Tumor | 26 | 0.269231 |
| CDK8 | chr13 c.606T>A | Yes | Tumor | 26 | 0.269231 |
| CDK8 | chr13 c.610C>T | Yes | Tumor | 26 | 0.269231 |
| CDK8 | chr13 c.615T>C | Yes | Tumor | 26 | 0.269231 |
| CDK8 | chr13 c.618T>C | Yes | Tumor | 26 | 0.269231 |
| CDK8 | chr13 c.624A>G | Yes | Tumor | 26 | 0.269231 |
| CDK8 | chr13 c.627G>A | Yes | Tumor | 26 | 0.269231 |
| FOXA1 | chr14 c.878G>T | Yes | Tumor | 10 | 0.2 |
| GNA13 | chr17 c.664G>C | Yes | Tumor | 139 | 0.438849 |
| **AKT2** | chr19 c.291G>A | Yes | Tumor | 69 | 0.173913 |
| **AKT2** | chr19 c.297G>A | Yes | Tumor | 70 | 0.171429 |
| **AKT2** | chr19 c.306C>T | Yes | Tumor | 73 | 0.164384 |
| **AKT2** | chr19 c.327C>T | Yes | Tumor | 82 | 0.146341 |
| **AKT2** | chr19 c.330C>G | Yes | Tumor | 83 | 0.144578 |
| **VAV1** | chr19 c.753A>G | Yes | Tumor | 16 | 0.125 |
| **SOS1** | chr2 c.653A>G | Yes | Tumor | 11 | 0.181818 |
| **GNAS** | chr20 c.125G>A | Yes | Tumor | 10 | 0.2 |
| **NCOA3** | chr20 c.90C>T | Yes | Tumor | 13 | 0.153846 |
| **FOXL2** | chr3 c.839C>T | Yes | Tumor | 15 | 0.133333 |
| **UBR5** | chr8 c.6714A>G | Yes | Tumor | 11 | 0.181818 |
| **MED12** | chrX c.5C>T | Yes | Tumor | 9 | 0.222222 |
| **HDAC7** | chr12 c.1620C>T | Yes | TC | 20 | 0.55 |
| **CD276** | chr15 c.471A>C | Yes | TC | 121 | 0.512397 |
| **CD276** | chr15 c.479C>T | Yes | TC | 118 | 0.5 |
| **SERPINB3** | chr18 c.836G>C | Yes | TC | 396 | 0.997475 |
| **INSR** | chr19 c.5C>G | Yes | TC | 5 | 1 |
| **ICOSLG** | chr21 c.1057T>C | Yes | TC | 68 | 1 |
| **DDX4** | chr5 c.993C>T | Yes | TC | 315 | 0.492063 |
| **H1-4** | chr6 c.455A>G | Yes | TC | 108 | 0.490741 |
| **MGAM** | chr7 c.2742A>C | Yes | TC | 214 | 0.537383 |
| **MGAM** | chr7 c.4820A>C | Yes | TC | 40 | 0.2 |
| **MGAM** | chr7 c.6389G>C | Yes | TC | 273 | 0.52381 |

**Table S10.** Table of cancer-associated genes with mutations that were exclusive to their respective organs in the CDX models. LN = Lymph node

| Liver Only |  | LN Only | Spleen Only |
| --- | --- | --- | --- |
| MTOR | PIK3CG | KDR | COP1 |
| DDR2 | MYC | RUNX1T1 | RHOA |
| SETDB1 | NTRK2 | RPS6KB2 | FOXL2 |
| RRAGC | SYK | NOTCH3 | EIF4A2 |
| MPL | ABL1 | EPOR | MGAM |
| JUN | GATA3 |  | LMO2 |
| JAK1 | SHOC2 |  | YAP1 |
| NOTCH2 | FGFR2 |  | HDAC7 |
| MCL1 | RET |  | IRS2 |
| PAX8 | NUP98 |  | YY1 |
| XPO1 | FGF3 |  | CD276 |
| IRS1 | RPS6KA4 |  | MAPK3 |
| CXCR4 | FLI1 |  | AURKB |
| PPARG | LRP5 |  | STAT5B |
| RAF1 | CCND2 |  | SOX9 |
| MST1R | NFE2 |  | AURKA |
| TP63 | ATF1 |  | BCR |
| MITF | MDM2 |  |  |
| FOXP1 | MSI1 |  |  |
| GATA2 | HIF1A |  |  |
| PIK3CA | SETD1A |  |  |
| DCUN1D1 | PPM1D |  |  |
| GAB1 | ALOX12B |  |  |
| PDGFRB | STAT5A |  |  |
| TLX3 | MAP3K14 |  |  |
| H1-4 | DNMT1 |  |  |
| ROS1 | INSR |  |  |
| H2AC17 | ERG |  |  |
| CARD11 | MED12 |  |  |
| ETV1 | BTK |  |  |

